# Mechanisms controlling membrane recruitment and activation of autoinhibited SHIP1

**DOI:** 10.1101/2023.04.30.538895

**Authors:** Grace L. Waddell, Emma E. Drew, Henry P. Rupp, Scott D. Hansen

## Abstract

Signal transduction downstream of growth factor and immune receptor activation relies on the production of phosphatidylinositol-(3,4,5)-trisphosphate (PI(3,4,5)P_3_) lipids by phosphoinositide-3-kinase (PI3K). Regulating the strength and duration of PI3K signaling in immune cells, Src homology 2 domain-containing inositol 5-phosphatase 1 (SHIP1) controls the dephosphorylation of PI(3,4,5)P_3_ to generate PI(3,4)P_2_. Although SHIP1 has been shown to regulate neutrophil chemotaxis, B-cell signaling, and cortical oscillations in mast cells, the role that lipid and protein interactions serve in controlling SHIP1 membrane recruitment and activity remains unclear. Using single molecule TIRF microscopy, we directly visualized membrane recruitment and activation of SHIP1 on supported lipid bilayers and the cellular plasma membrane. We find that SHIP1’s interactions with lipids are insensitive to dynamic changes in PI(3,4,5)P_3_ both in vitro and in vivo. Very transient SHIP1 membrane interactions were detected only when membranes contained a combination of phosphatidylserine (PS) and PI(3,4,5)P_3_ lipids. Molecular dissection reveals that SHIP1 is autoinhibited with the N-terminal SH2 domain playing a critical role in suppressing phosphatase activity. Robust SHIP1 membrane localization and relief of autoinhibition can be achieved through interactions with immunoreceptor derived phosphopeptides presented either in solution or conjugated to supported membranes. Overall, this work provides new mechanistic details concerning the dynamic interplay between lipid binding specificity, protein-protein interactions, and activation of autoinhibited SHIP1.

## INTRODUCTION

Phosphatidylinositol phosphate (PIP) lipids play a crucial function in eukaryotic cell biology by regulating the localization and activity of numerous signaling proteins on intracellular membranes (1). PIP lipids are transiently created by different classes of lipid kinases and phosphatases, which are activated during various cell signaling pathways (2). Understanding how PIP lipid modifying enzymes are regulated is critical for determining how cells control the strength and duration and PIP lipid signaling in cell biology. Misregulation of PIP lipid homeostasis has been shown to profoundly affect cell growth and proliferation, which is linked to poor prognosis in numerous human diseases (3).

The Src homology 2 domain-containing inositol 5-phosphatase 1 (SHIP1) is a hematopoietic cell specific lipid phosphatase that dephosphorylates phosphatidylinositol-(3,4,5)-trisphosphate (PI(3,4,5) P_3_) to generate phosphatidylinositol-(3,4)-bisphosphate (PI(3,4)P_2_). PI(3,4,5)P_3_ is a second messenger lipid that selectively localizes a myriad of signaling proteins (e.g. Akt, BTK, P-Rex1) to the plasma membrane that are critical for cell growth, survival, and development (4–6). Because PI(3,4,5)P_3_ lipids regulate critical cellular processes, these lipids need to be spatially and temporally regulated for cellular homeostasis. Cells lacking SHIP1 display defects in cell adhesion and migration as a consequence of elevated PI(3,4,5)P_3_ levels (7–9).

To access its lipid substrate, SHIP1 must localize to the plasma membrane. Antibody induced activation of the immunoglobulin E receptor (FcεRI) in mast cells revealed that SHIP1 undergoes dynamic cycles of plasma membrane recruitment and dissociation which leads to the emergent property of cortical oscillations (10). It was also shown that stimulating the B cell- and FcγRIIB-receptors enhances SHIP1 plasma membrane localization compared to unstimulated cells (11, 12). Although SHIP1 is recruited to the plasma membrane upon activation of several types of immune cell receptors, the mechanism controlling membrane recruitment and activation is unclear. In particular, it’s unclear whether SHIP1 is recruited to the plasma membrane through direct interactions with PI(3,4,5)P_3_ lipids or indirectly by proteins that bind to PI(3,4,5)P_3_. Understanding how each domain of SHIP1 contributes to membrane localization and lipid phosphatase activity will close the gap in knowledge concerning whether SHIP1 directly interacts with activated receptors, newly synthesized PIP lipids, or proteins that bridge these interactions.

SHIP1 contains two proposed lipid binding domains that flank the central phosphatase domain and have been proposed to regulate membrane localization and catalytic activity (**Figure 1A**). The pleckstrin-homology related (PH-R) domain was shown to interact with the substrate of SHIP1, PI(3,4,5)P_3_, while the C2 domain reportedly binds to the product of SHIP1, PI(3,4)P_2_ (13–15). Mutations in the PH-R domain disrupted localization to the phagocytic cup during FcγR-mediated phagocytosis in macrophages but did not affect SHIP1 catalysis. This suggests that SHIP1 localization to PI(3,4,5)P_3_ containing membranes is independent of its catalytic domain (13). Biochemical analysis of the SHIP1 paralog, SHIP2, revealed a structural interface between the phosphatase and C2 domain capable of interdomain communication and allosteric regulation (16). The C2 domain of SHIP2 reportedly interacts with phosphatidylserine (PS) (16), while the SHIP1 C2 domain reportedly binds to PI(3,4)P_2_ (14). Based on sequence homology between SHIP1 and SHIP2, both phosphatases are predicted to have similar phospholipid binding specificity (see Supplemental Information). Together, these reported lipid interactions suggest a possible mechanism for positive feedback and allosteric activation during SHIP1-mediated dephosphorylation of PI(3,4,5)P_3_.

**Figure 1.**
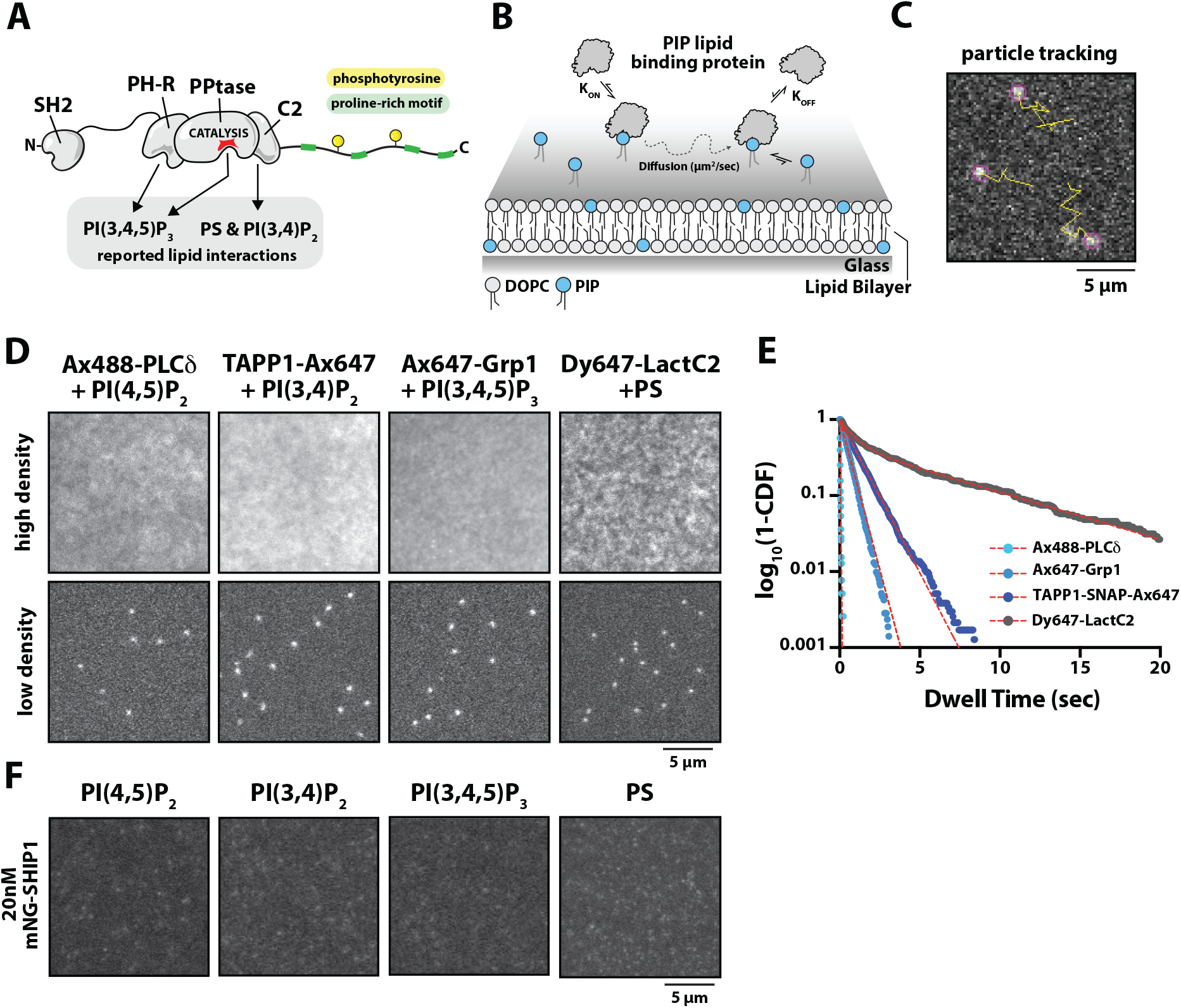
SHIP1(PH-PP-C2) does exhibit strong interactions with individual PIP lipids in vitro. **(A)** Cartoon diagram showing the reported lipid binding domains of SHIP1, which include the phosphatase domain and flanking PH-R and C2 domains. **(B)** Experimental setup for directly visualizing protein-lipid interactions on supported lipid bilayers using smTIRF-M. **(C)** Representative image showing single particle detection (purple circle) and trajectories (yellow line) of 3 Ax647-Grp1 membrane bound molecules. Image collected in the presence of 1 pM Ax647-Grp1. Membrane composition: 2% PI(3,4,5)P_3_ and 98% DOPC. **(D)** Representative TIRF-M images showing bulk membrane recruitment in the presence of either 20 nM (“high density”) or (“low density”) Ax488-PLCδ (250 pM), TAPP1-SNAP-Ax647 (1 pM), Ax647-Grp1 (1 pM), or LactC2-Dy647 (2 pM) on SLBs containing DOPC lipids plus either 2% PI(4,5)P_2_, 2% PI(3,4)P_2_, 2% PI(3,4,5)P_3_, or 20% DOPS lipids, respectively. **(E)** Single molecule dwell time distributions of various PIP lipid binding domains plotted as dwell time versus log_10_(1-cumulative distribution frequency). Curves are fit with a single or double exponential decay curve yielding the following dwell times: Ax488-PLCδ (τ_1_ = 24 ± 2 ms) TAPP1-SNAP-Ax647 (τ_1_ = 1.02 ± 0.053 s), Ax647-Grp1 (τ_1_ = 0.544 ± 0.007 s), or LactC2-Dy647 (τ_1_ = 0.765 ± 0.191 s, τ_2_ = 6.58 ± 0.539 s, α = 0.5 ± 0.04). See *Table S1* for statistics. **(F)** mNG-SHIP1(PH-PP-C2) does not robustly associated with SLBs containing PIP or PS lipids. Representative images showing bulk membrane recruitment in the presence of 20 nM mNG-SHIP1(PH-PP-C2) on SLBs containing DOPC lipids plus either 2% PI(4,5)P_2_, 2% PI(3,4)P_2_, 2% PI(3,4,5)P_3_, or 20% DOPS, respectively. Note that image intensities were scaled in a similar manner to the top row of images shown in (C).

Previous biochemical characterization of SHIP1 suggests that the C-terminal domain attenuates 5-phosphatase activity. In particular, a C-terminal truncation was previously shown to increase SHIP1 activity approximately 8-fold compared to full-length (FL) SHIP1 (17). Consistent with this result, C-terminal truncation mutants of SHIP2 were also more active than the full-length phosphatase (18). A potential mechanism for autoinhibition could be mediated through intramolecular interactions between the N-terminal SH2 domain and C-terminal phosphotyrosine(s). Supporting this model, a peptide derived from the C-terminal region of SHIP1 containing phosphorylated tyrosine residue Y1022 was previously shown to be immunoprecipitated purified SH2 domains of SHIP1 (19). Similar to Src-family tyrosine kinases (20, 21), the N-terminal SH2 domain of SHIP1 could potentially form an intramolecular auto-inhibitory interaction with a C-terminal phosphotyrosine residue (e.g. Y1022). Although Y1022 was shown to be phosphorylated by Src family kinases (22), it remains unclear whether this post-translational modification is responsible for SHIP1 autoinhibition or if it regulates interactions with peripheral membrane binding proteins. Despite evidence supporting SHIP1 autoinhibition, the mechanism that regulates the formation and relief of intramolecular interactions remains unknown. Understanding the mechanism of autoinhibition could reveal how SHIP1 is recruited to the plasma membrane and the functional consequences of SHIP1 mutants lacking autoinhibitory contacts.

To date, no biochemical studies have directly visualized the interactions between purified SHIP1 and proposed PIP lipid binding partners. Previous experiments have relied on surface plasma resonance (SPR) (15) and protein lipid overlay (PLO) assays (Ong et al. 2007, Ming-Lum et al. 2012) to measure SHIP1 lipid interactions. Given the combination of protein and lipid interactions that potentially regulate SHIP1, it remains unclear whether SHIP1’s membrane localization is controlled directly or indirectly by PIP lipid interactions. To directly visualize proposed interactions between SHIP1 and different PIP lipid species, we established a single molecule Total Internal Reflection Fluorescence (smTIRF) Microscopy assay in combination with supported lipid bilayers (SLBs). Direct visualization of fluorescently labeled SHIP1 revealed very weak lipid binding specificity compared to several well-established PIP lipid binding proteins that display a range of affinities (i.e. K_D_ = 0.01 – 10 μM) (23–27). Similarly, in live cells, we find that central lipid phosphatase domain of SHIP1 flanked by the two proposed lipid binding domains transiently associates with the plasma membrane and is non-responsive to G-protein coupled receptor (GPCR) induced changes in PI(3,4,5)P_3_ levels. Reconstitution of SHIP1 lipid interactions in vitro on SLBs revealed that a combination of phosphatidylserine (PS) and PI(3,4,5)P_3_ lipids was required to detect the lower limit of transient membrane dwell times. Biochemical reconstitution of SHIP1 phosphatase activity revealed that the full-length enzyme is autoinhibited compared to the central PH-PP-C2 domains. Characterization of SHIP1 truncation mutants indicates that both the N-terminal SH2 domain and the disordered C-terminal tail limit SHIP1 phosphatase activity, but to different extents. In the presence of an immune receptor derived phosphopeptide presented either in solution or membrane tethered, SHIP1 lipid phosphatase activity can be robustly stimulated. Overall, our results reveal that interactions between SHIP1 and various lipid species are surprisingly weak and likely serve a secondary role following membrane recruitment mediated by SHIP1-protein interactions.

## RESULTS

### SHIP1 displays low specificity for proposed PIP lipid binding partners

Flanking the central phosphatase domain, SHIP1 contains two proposed lipid binding domains (**Figure 1A**). The pleckstrin-homology related domain (herein referred to as “PH”) was previously shown to interact with the substrate of SHIP1, PI(3,4,5)P_3_ (13), while the C2 domain reportedly binds to SHIP1’s product, PI(3,4)P_2_ (14). These lipid interactions are hypothesized to function in concert to regulate SHIP1 membrane docking and lipid phosphatase activity. A major limitation in our understanding of these lipid interactions is that previous experiments utilized binding assays that did not directly visualize dynamic membrane association, bilayer diffusion, and dissociation of SHIP1 (**Figure 1B**). Observing these behaviors are essential criteria needed to validate that proposed lipid binding domains autonomously and reversibly interact with specific lipid species.

To assess the role that lipids serve in controlling SHIP1 membrane localization, we established a method to directly visualize the dynamic and reversible membrane binding interactions using supported lipid bilayers (SLBs) visualized by Total Internal Reflection Fluorescence (TIRF) microscopy (**Figure 1B-1C**). We purified and fluorescently labeled a collection of lipid binding domains derived from Lact-C2, PLCδ, TAPP1, and Grp1. These domains have well established interactions with either phosphatidylserine (PS), PI(4,5)P_2_, PI(3,4)P_2_, or PI(3,4,5)P_3_ respectively (23–27). Each protein has a K_D_ in the range of 0.01 – 10 μM, providing a panel of positive controls for directly visualizing various PIP lipid interactions in vitro. When we measured the localization of these lipid binding domains on SLBs using a solution concentration of 20 nM for each PIP lipid binding protein, we observed robust membrane recruitment based on the high membrane fluorescence intensity that blankets the SLB compared to the background signal (**Figure 1D**). To compare the relative strengths of these protein-lipid interactions, we measured the dwell time of single spatially resolved membrane binding events (**Figure 1E, Movie S1-S4**). Consistent with previous K_D_ measurements based on solution binding assays, we calculated a range of single molecule dwell times (t_1_ = 0.02 – 5 seconds) for these lipid binding proteins (**Figure 1E, Figure S1, Table S1**).

In order to measure single molecule interactions between SHIP1 and various phospholipids, we recombinantly expressed and purified a mNeonGreen (mNG) tagged SHIP1 truncation containing the central phosphatase domain flanked by two lipid binding domains (i.e. mNG-PH-PP-C2). Compared to the membrane localization of the established lipid binding domains (**Figure 1D-1E**), we observed no significant membrane association of mNG-SHIP1(PH-PP-C2) on SLBs containing either PS, PI(4,5)P_2_, PI(3,4)P_2_, or PI(3,4,5)P_3_ lipids alone (**Figure 1F**). Although a small fraction of molecules bound to the supported lipid bilayer, the lack of diffusivity indicated that those binding events represent non-specific interactions with diffraction limited defects in the supported membrane. Increasing the frame rate of image acquisition to >80 frames per seconds, allowed us to observe low frequency membrane binding events that lasted only 1 frame (12 ms) in the presence of either PS or PI(3,4,5)P_3_ lipids alone. Overall, the mNG-SHIP1(PH-PP-C2) membrane interactions were too transient for us to calculate a characteristic dwell time using any of the membrane compositions described above.

### Plasma membrane localization of SHIP1(PH-PPtase-C2) is transient and insensitive to dynamic PI(3,4,5)P_3_ production

Our *in vitro* characterization of SHIP1 on supported membranes by smTIRF lacks some of the complexity that exists on the cellular plasma membrane. To determine whether the membrane binding behavior of SHIP1(PH-PP-C2) is significantly different *in vitro* and *in vivo*, we established conditions to visualize the plasma membrane localization of PIP lipid binding domains (i.e. Grp1 and Lact-C2) and SHIP1(PH-PP-C2) in neutrophil-like PLB-985 cells with single molecule resolution. For these experiments, we tagged Grp1, LactC2, and SHIP1 with a green to red photoconvertible mEos3.2 (referred to as mEos) (**Figure 2A**). Genes encoding these proteins were transduced into PLB-985 cells using lentivirus, which resulted in a uniform localization pattern across the plasma membrane (**Figure 2B**). To resolve single membrane binding events, we photoconverted a fraction of the mEos tagged proteins to the more photostable red fluorescent state using a transient pulse of 405 nm light (**Figure 2B**). The single molecule brightness distribution and photobleaching analysis indicate that tracked mEos-tagged molecules represent single spatially resolved fluorescent proteins (**Figure 2C-2D**). Single molecule dwell time analysis of plasma membrane localized mEos-Grp1 and mEos-LactC2 produced very similar dwell times of 392 ms and 371 ms, respectively (**Figure 2E, Movie S5-S6, Table S1**). Given that the dwell times of fluorescently labeled Grp1 and LactC2 were longer in our in vitro SLB assay, the in vivo dwell times for these lipid binding proteins likely represents the upper limit for single molecule track length measured in cells under our experimental conditions. By comparison mEos-SHIP1(PH-PP-C2) expressed in PLB-985 cells exhibited plasma membrane interactions that were comparatively transient in nature (τ_1_ = 38 ms) (**Figure 2I, Movie S7, Table S1**). Notably in live cells, we observed a large fraction of photoconverted mEos-SHIP1(PH-PP-C2) rapidly moving in and out of the TIRF-M focal plane (**Movie S7**). This population of molecules appears as a diffuse fluorescent cloud near the plasma membrane and indicates that the vast majority of SHIP1 molecules localizes to the plasma membrane, but are unable to persistently engage lipids to produce a spatially resolved and quantifiable dwell time.

**Figure 2.**
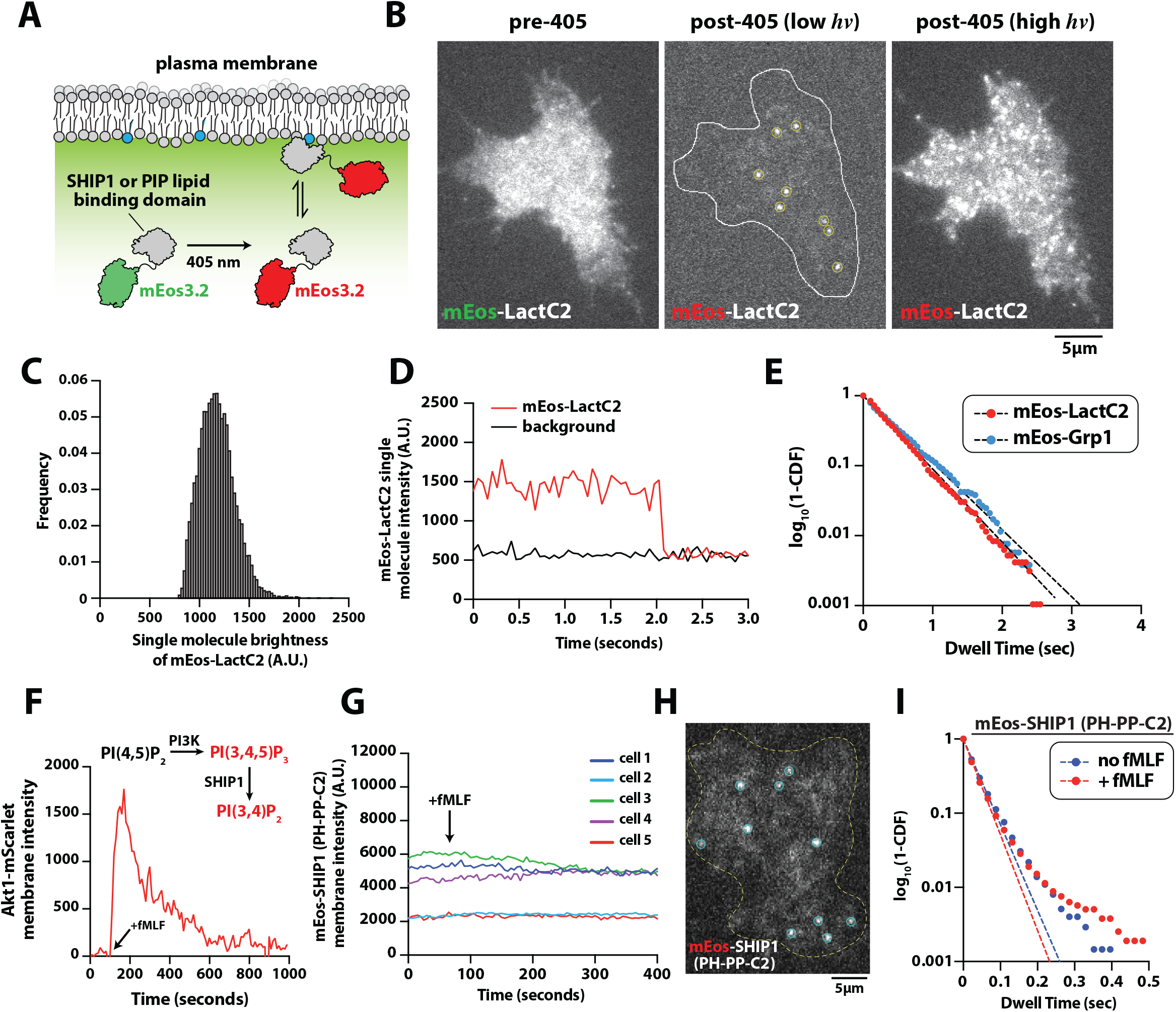
SHIP1 (PH-PP-C2) plasma membrane localization is insensitive to dynamic changes in PI(3,4,5)P_3_ lipid composition. **(A)** Cartoon diagramming UV-dependent photoconversion and plasma membrane binding of a mEos tagged cytoplasmic protein in cells. **(B)** Representative images showing localization of mEos-LactC2 before and after photoconversion with 405 nm UV light (“low” and “high” laser power). Localization of mEos-LactC2 was visualized using smTIRF-M in PLB-985 cells. **(C)** Molecular brightness distribution of single plasma membrane localized mEos-Lact-C2 molecules. **(D)** Stepwise photo-bleaching of a single plasma membrane localized mEos-Lact-C2 molecules compared to the background fluorescence of the cell. **(E)** Single molecule dwell distributions of mEos-Lact-C2 and mEos3.2-Grp1. Curves were fit to a single exponential decay curve to calculate the following dwell times: mEos-Lact-C2 (τ_1_ = 371 ± 7 ms, N=12 cells) and mEos-Grp1 (τ_1_ = 392 ± 5 ms, N=10 cells). **(F)** Plot showing the translocation of Akt1-mScarlet to the plasma membrane following stimulation of PLB-985 cells with 10 nM fMLF. **(G)** Bulk membrane localization of non-photoconverted mEos-SHIP1(PH-PP-C2) does not change in response to stimulating PLB-985 cells with 10 nM fMLF (arrow). Traces represent the single cell plasma membrane intensity of mEos-SHIP1(PH-PP-C2) measured by TIRF-M. **(H)** Representative image showing single particle detection of mNG-SHIP1 in live PLB-985 cells. **(I)** Single molecule dwell time distributions of photoconverted mEos-SHIP1(PH-PP-C2) in PLB-985 cells ± 10 nM fMLF plotted as dwell time versus log_10_(1-cumulative distribution frequency). Curves are fit with a single exponential decay curves and both curves yield the following dwell time for mEos-SHIP1(PH-PP-C2): τ_1_ = 38 ± 3 ms (pre-fMLF; N=4 cells) and τ_1_ = 37 ± 5 ms (post-fMLF; N=5 cells). A single dwell time was measured for each cell with n = 421-4267 molecules tracked per cell. Dwell times are reported as mean ± SD.

To determine whether the membrane binding avidity of mEos-SHIP1(PH-PP-C2) could be modulated by dynamic changes in substrate or product availability at the plasma membrane, we stimulated PLB-985 neutrophil-like cells with a uniform concentration of chemoattractant, N-formyl-methionine-leucine-phenylalanine (fMLF). Activation of the formyl peptide receptor with fMLF results in a transient spike in PI(3,4,5)P_3_ and PI(3,4)P_2_ production (28, 29), which we visualized using the dual specificity lipid binding domain, Akt1-mScarlet (**Figure 2F**). Following PLB-985 cell activation with 10 nM fMLF, we observed no change in the bulk plasma membrane localization of mNG-SHIP1(PH-PP-C2) (**Figure 2G**). Similarly, single molecule dwell time analysis revealed that membrane avidity of mEos-SHIP1(PH-PP-C2) was insensitive to dynamic changes in PI(3,4,5)P_3_ and PI(3,4)P_2_ levels (**Figure 2I, Table S1, Movie S7**). Consistent with our in vitro observations, this result supports an absence of high affinity interactions between SHIP1(PH-PP-C2) and either PI(3,4,5)P_3_ or PI(3,4)P_2_ lipids.

### SHIP1 catalyzes dephosphorylation of PI(3,4,5)P_3_ with unimolecular kinetics

Based on reported lipid interactions, researchers have hypothesized that SHIP1 catalyzes the dephosphorylation of PI(3,4,5)P_3_ with a positive feedback loop based on interactions with both substrate and product (13–15). To measure potentially nonlinear reaction kinetics, we developed an in vitro assay to simultaneously visualize SHIP1 membrane binding and lipid phosphatase activity using TIRF microscopy. For these experiments, we used both fluorescently labeled Grp1 and TAPP1 PH domains to monitor changes in the membrane abundance of PI(3,4,5)P_3_ and PI(3,4)P_2_ (**Figure S2A-S2C**). Using fluorescently labeled TAPP1, we found that the interaction with PI(3,4)P_2_ was quite strong resulting in continual membrane binding even after SHIP1-mediated depletion of PI(3,4,5)P_3_. This made quantification of reaction completion times ambiguous under experimental conditions that utilize varying concentrations of PS lipids concentration. For this reason, we primarily used the Alexa647-Grp1 PH domaintomonitor SHIP1-mediateddephosphorylation of PI(3,4,5)P_3_ on supported membranes (**Figure 3A, Figure S2D-S2F**). To display kinetic traces in terms of product formation, we inverted the Grp1 membrane dissociation curves to represent the time dependent increase in SHIP1 product formation (**Figure 3B**). Independent of how data was plotted, kinetic traces were hyperbolic shaped (**Figure 3B, Figure S2**). Consistent with SHIP1 catalyzing reactions with unimolecular kinetics, we observed no reciprocal regulation linking changes in PI(3,4,5)P_3_ or PI(3,4)P_2_ abundance to membrane localization of SHIP1 (**Figure 3C**). If there was any reciprocal regulation resulting from lipid interactions, albeit weak, SHIP1 membrane recruitment curves would be more sigmoidal in shape and increase with production of PI(3,4)P_2_ lipids. However, membrane fluorescent intensity of mNG-SHIP1(PH-PP-C2) remained constant throughout the course of the reaction (**Figure 3C**). This was consistent with our inability to detect interactions between mNG-SHIP1(PH-PP-C2) and either PI(3,4,5) P_3_ or PI(3,4)P_2_ lipids by smTIRF (**Figure 1E-1F**).

**Figure 3.**
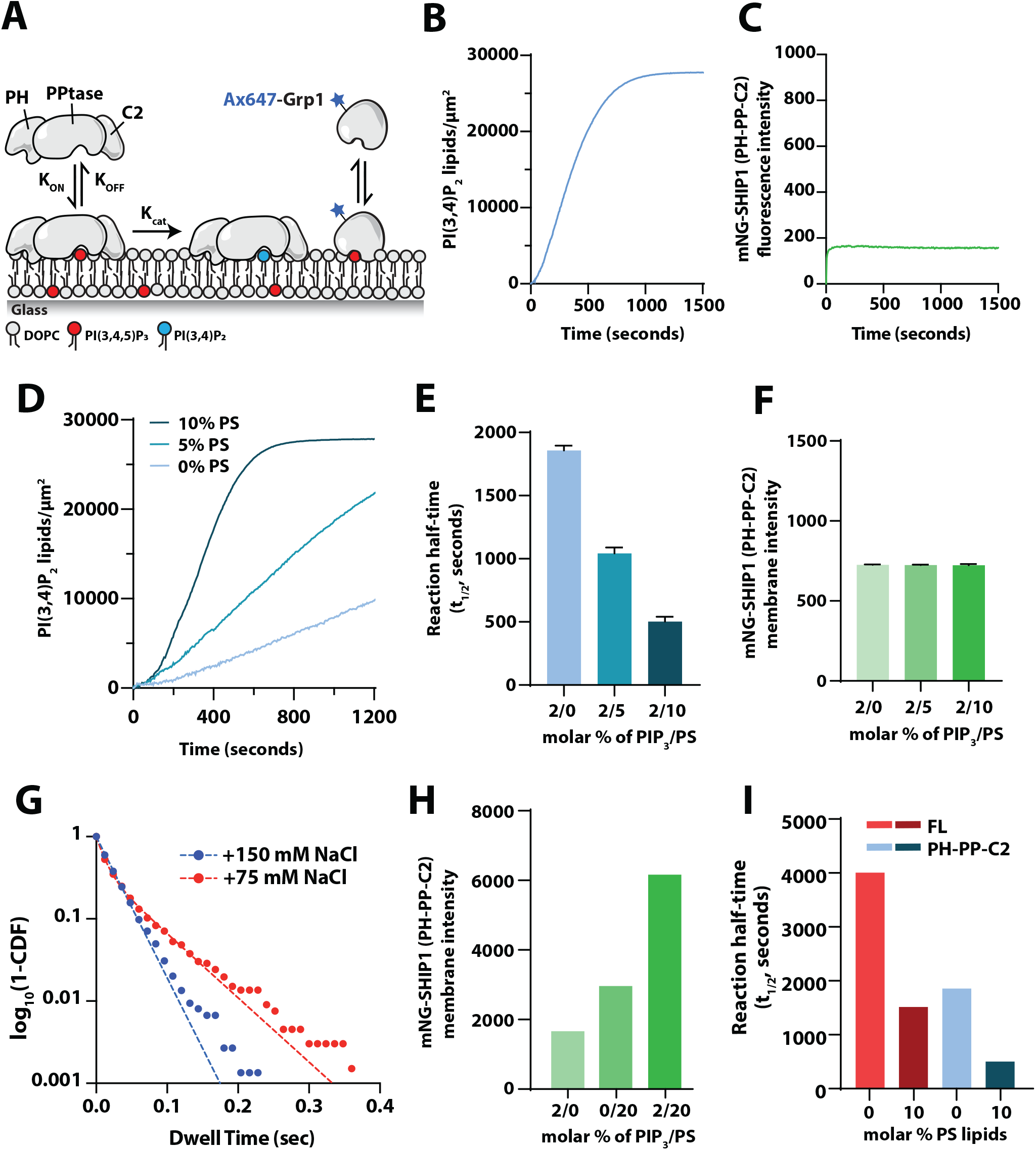
SHIP1 catalyzes the dephosphorylation of PI(3,4,5)P_3_ with simple kinetics. **(A)** Experimental design for measuring SHIP1 lipid phosphatase activity in vitro on supported lipid bilayers using TIRF-M. **(B)** Kinetics of 20nM SHIP1(PH-PP-C2). PI(3,4,5)P_3_ dephosphorylation measured using 20 nM Ax647-Grp1. **(C)** Bulk membrane localization of 20nM mNG-SHIP1 during catalysis shown in (B). **(D)** PS lipids enhance SHIP1 phosphatase activity. Representative kinetics traces of 4 nM mNG-SHIP1 (PH-PPtase-C2 domain) in the absence and presence of PS lipids. Production of PI(3,4)P_2_ was monitored by the presence of 20 nM Ax647-Grp1. Initial membrane composition: 88-98% DOPC, 0-10% PS lipids, and 2% PI(3,4,5)P_3_. **(E)** Quantification of reaction half time in (D). Bars equal mean values [N = 2 reactions per concentration, error = SD]. **(F)** Quantification of bulk localization measured in the presence of 20 nM mNG-SHIP1(PH-PPtase-C2) on SLBs containing 2% PI(3,4,5)P_3_ and 0, 5, 10% PS lipids [N = 10 fluorescent intensity measurements per membrane, error = SD]. **(G)** Single molecule dwell time measured in the presence of 200 pM mNG-SHIP1(PH-PPtase-C2). Data plotted as dwell time versus log_10_(1-cumulative distribution frequency). Curves fit with a single or double exponential decay curve yielding the following dwell times: 150 mM NaCl buffer (τ_1_ = 25 ± 1 ms) or 75 mM NaCl buffer (τ_1_ = 9 ± 2 ms, τ_2_ = 56 ± 7 ms, α = 0.44 ± 0.13). Membrane composition: 88% DOPC, 2% PI(3,4,5)P_3_, 20% PS. **(H)** Quantification of bulk localization measured in the presence of 20 nM mNG-SHIP1(PH-PPtase-C2) on SLBs containing 2% PI(3,4,5)P_3_ and 0, 5, 10% PS lipids. **(I)** Full-length SHIP1 is stimulated by PS lipids, but exhibits lower activity compared to mNG-SHIP1(PH-PPtase-C2). Reaction half times comparing phosphatase activity in the presence of 4nM full-length mNG-SHIP1 or 4nM mNG-SHIP1(PH-PPtase-C2). Membrane composition: 88-98% DOPC, 2% PI(3,4,5)P_3_, +/-10% PS..

Phosphatidylserine (PS) lipids have been reported to bind to the C2 domain of SHIP2 and allosterically regulate its phosphatase activity (16). Consistent with this model, we measured enhanced SHIP1 phosphatase activity over a range of PS lipid densities (**Figure 3D-3E**). However, the increase in activity was not correlated with a dramatic change in bulk membrane localization of mNG-SHIP1(PH-PP-C2) (**Figure 3F**). PS dependent enhancements in activity without any associated enhancement in bulk membrane localization was surprising, but consistent with a mechanism of allostery or regulation of the SHIP1 membrane docking orientation. Next, we returned to single molecule dwell time analysis of mNG-SHIP1(PH-PP-C2) using supported membranes containing a mixture of 2% PI(3,4,5)P_3_ and 20% PS lipids. Under these conditions, we could detect very transient mNG-SHIP1(PH-PP-C2) membrane binding dynamics (τ_1_ = 25 ms, **Figure 3G, Table S1, Movie S8**), similar to our in vivo observations (**Figure 2I**). To strengthen the weak interactions observed between mNG-SHIP1(PH-PP-C2), PI(3,4,5)P_3_, and PS lipids, we reduced the buffer ionic strength (75 mM NaCl) and repeated out dwell time analysis. Under these conditions, membrane associated mNG-SHIP1(PH-PP-C2) changed to display two characteristic dwell times (τ_1_ = 9 ms, τ_2_ = 56 ms, α= 44%, **Figure 3G, Table S1**). In the presence of lower ionic strength buffer, we also observed a modest increase in bulk membrane localization of mNG-SHIP1(PH-PP-C2) that was dependent on PI(3,4,5)P_3_ and PS lipids (**Figure 3H**).

Next, we aimed to determine if PS lipids could similarly increase phosphatase activity of full-length SHIP1 (1-1188aa) (FL SHIP1). Although we found that PS lipids enhanced the activity of FL SHIP1, the overall active was still reduced to SHIP1(PH-PP-C2) (**Figure 3I**). Although PS lipids are considered allosteric regulators of SHIP1, this result suggested that the domains flanking the PH-PP-C2 domains suppress the catalytic activity of SHIP1.

### Full-length SHIP1 is autoinhibited by the N-terminal SH2 domain

It was previously demonstrated that truncating the C-terminus of SHIP1 enhances phosphatase activity of FL SHIP1, suggesting the C-terminus negatively regulates phosphatase activity (17). By contrast, studies using SHIP2 immunoprecipitated from cell lysate found that the N-terminal SH2 domain in conjunction with the C-terminus inhibits SHIP2 activity (18). To investigate potential mechanisms of SHIP1 autoinhibition, we purified full-length (FL) recombinant SHIP1, an N-terminal truncation mutant (ΔSH2), and a C-terminal truncation mutant (ΔCT) using baculovirus expression in insect cells (**Figure 4A**). Using our SLB TIRF assay, we measured the phosphatase activity of FL SHIP1 and truncation mutants. These experiments revealed that FL SHIP1 has reduced activity compared to SHIP1 (PH-PP-C2) construct (**Figure 4B**). Although it was previously demonstrated that truncating the C-terminus of SHIP1 increased activity 9-fold (17), removal of this region only increased SHIP1 activity 1.8 fold compared to the FL protein (**Figure 4B-4D**). The C-terminus of SHIP1 has two NPxY motifs that can be phosphorylated by Src family kinases. However, the increase in activity observed for the ΔCT mutant is unlikely a result of post translational modification. Incubation of SHIP1 (ΔCT) with the promiscuous tyrosine phosphatase, YopH, did not modulate phosphatase activity (**Figure S3**).

**Figure 4.**
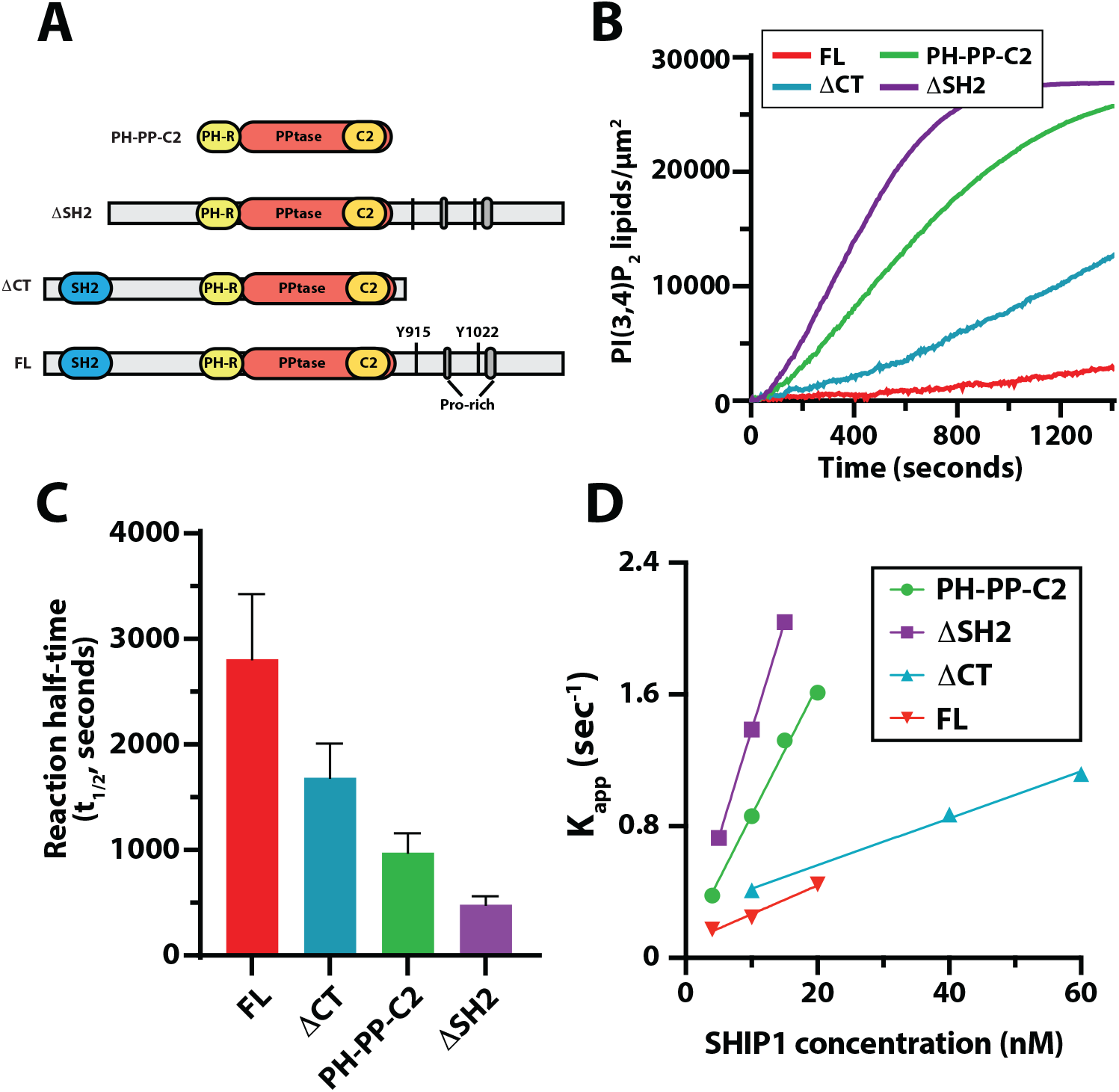
Full-length SHIP1 is autoinhibited by the N-terminal SH2 domain. **(A)** Domain organization of full-length (FL) SHIP1 and truncation mutants used in supported lipid bilayer TIRF-M assays. **(B)** Kinetic traces showing the dephosphorylation of PI(3,4,5)P_3_ in the presence 10 nM mNG tagged FL SHIP1, SHIP1(PH-PPtase-C2), SHIP1(ΔSH2), and SHIP1(ΔCT). **(C)** Quantification of reaction half times measured in the presence of 10 nM SHIP1, FL and truncation mutants. Bars equal mean values [N = 4-5 technical replicates, error = SD]. **(D)** Plot shows the relationship between SHIP1 concentration and phosphatase activity. comparing FL SHIP1, SHIP1(PH-PPtase-C2), SHIP1 (ΔSH2), and SHIP1 (ΔC-terminus). **(B-F)** Initial membrane composition: 98% DOPC, 2% PI(3,4,5)P_3_.

Biochemical characterization of the SHIP2 isoform found that the N-terminal SH2 domain in conjunction with the C-terminus inhibits SHIP2 activity (18). In our experiments, removal of the N-terminal SH2 domain of SHIP1 alone yielded an activity curve nearly identical to our PH-PP-C2 truncation mutant, suggesting that SHIP1 autoinhibition is largely dependent on the SH2 domain (**Figure 4B-4D**).

### Membrane recruitment and activation of SHIP1 by a phosphotyrosine peptide

During cell signaling, Src-Homology 2 (SH2) domains play a critical role regulating membrane recruitment of cytoplasmic proteins to immune receptors that contain tyrosine phosphorylated motifs (pY) (22, 30). Because truncating the SH2 domain of SHIP1 relieved autoinhibition, we hypothesized that phosphopeptide interactions with the SH2 domain could activate SHIP1 (**Figure 5A**). To decipher this mechanism, we derived a peptide from the immunoreceptor tyrosine inhibitory motif (ITIM) of the FcγRIIB receptor containing a single phosphotyrosine residue (pY-ITIM). SHIP1 is reported to be critical for the inhibitory function of FcγRIIB (31), and this peptide was previously shown to interact with the SH2 domain of SHIP1 in a pulldown assay (19). In solution, the pY-ITIM peptide enhanced the activity of FL SHIP1 (**Figure 5B-5C**).

**Figure 5.**
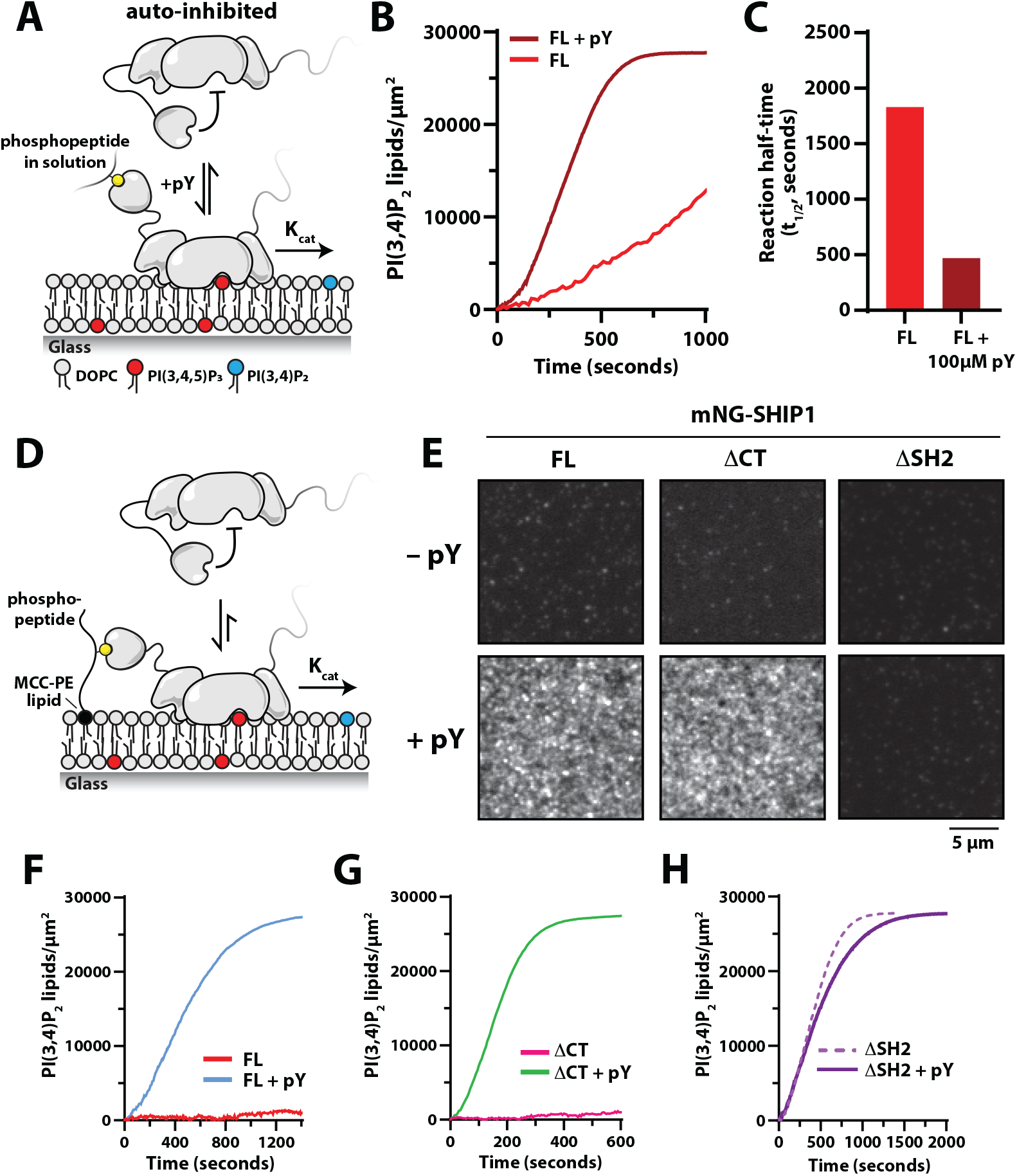
Phosphotyrosine peptides promote membrane recruitment and activation of SHIP1. **(A)** Experimental design for measuring SHIP1 phosphatase activity in the presence phosphotyrosine peptides derived from ITIM motif of the FcγRIIB receptor (pY-ITIM) in solution. **(B)** 100 μM pY-ITIM in solution stimulates the phosphatase activity of 20nM full-length SHIP. Initial membrane composition: 96% DOPC, 2% PI(3,4,5)P_3_, 2% MCC-PE (quenched/non-reactive). **(C)** Quantification of reaction half times in (B). **(D)** Experimental design for measuring SHIP1 phosphatase activity in the presence of membrane tethered pY-ITIM peptides. **(E)** Representative TIRF-M images showing localization of 5 nM mNG-SHIP1(FL, 1-1188aa), mNG-SHIP1(ΔCT), and mNG-SHIP1(ΔSH2) in the presence of pY-ITIM conjugated to the membrane. **(F)** Membrane conjugated pY-ITIM peptide stimulates 0.1 nM SHIP1 (FL) phosphatase activity. **(G)** Membrane conjugated pY-ITIM peptide stimulates 0.1 nM SHIP1 (ΔCT) phosphatase activity. **(H)** Membrane conjugated pY-ITIM peptide does not stimulates 5 nM SHIP1 (ΔSH2) phosphatase activity. **(F-H)** Plots show representative kinetics traces. Production of PI(3,4)P_2_ was monitored by the presence of 20 nM Ax647-Grp1. Plots are inverted to show production of PI(3,4)P_2_. **(E-H)** Membrane composition: 96% DOPC, 2% PI(3,4,5)P_3_, 2% MCC-PE-(pY conjugated).

*In vivo*, SHIP1 is recruited to the plasma membrane by pY-ITIM following receptor activation in B-cells (22). To determine how membrane anchored pY-ITIM regulates SHIP1 membrane recruitment and activation, we covalently attached the pY-ITIM peptide to a cysteine reactive maleimide lipid that was incorporated into supported membranes containing PI(3,4,5)P_3_ lipids (**Figure 5D**). We found that pY-ITIM covalently coupled to lipids rapidly recruited and activated SHIP1 compared to experiments performed with the pY-ITIM peptide in solution (**Figure 5E-5F**). We also measured rapid activation of the partially autoinhibited C-terminal truncation mutant (ΔCT) and localization to SLBs (**Figure 5E, 5G**). By contrast, we did not observe pY dependent localization of the N-terminal truncation mutant (ΔSH2) (**Figure 5H**), supported by the absence of pY binding (**Figure 5E**). Together, these results further support an autoinhibitory mechanism that’s mediated by the N-terminal SH2 domain and demonstrates that receptor derived ITIMs can both membrane recruit and stimulate SHIP1 activity.

## DISCUSSION

### SHIP1 lipid binding specificity

SHIP1 is a multidomain protein with two proposed lipid binding domains considered important for membrane localization and activity (13, 14) (**Figure 1A**). Due to a lack of structural data and mutational analysis of these domains, it has been difficult to define the molecular basis of proposed SHIP1 lipid interactions. To gain insight about the nature of SHIP1 lipid binding specificity, we directly visualized membrane association and dissociation dynamics of fluorescently labeled SHIP1 of supported lipid bilayers (SLBs) using single molecule TIRF-M (smTIRF-M). Despite reports of PIP lipid interactions being mediated by both the pleckstrin-homology related (PH-R) (13) or C2 domains (14), our *in vitro* single molecule dwell time analysis indicates that SHIP1 does not strongly associate with PI(4,5)P_2_, PI(3,4)P_2_, PI(3,4,5)P_3_, or PS lipids that are incorporated into supported membranes (**Figure 1F**). Based on the reported affinity that SHIP1 has for PI(3,4)P_2_ (*K*_*D*_ = 6 nM) and PI(3,4,5)P_3_ (*K*_*D*_ = 1 nM) (15), we expected to observe single molecule dwell times greater than 10 seconds (*K*_*OFF*_ = 0.1 sec^-1^). This estimate assumes that SHIP1 membrane association is diffusion limited (*K*_*ON*_ = 10 μM^-1^sec^-1^). Based on the proposed reciprocal regulation between SHIP1 and PIP lipids interactions, we also expected SHIP1 localization to be cooperative in nature and for activity traces to display positive feedback based on product recognition. Instead, SHIP1 catalyzed PI(3,4,5)P_3_ dephosphorylation with binding interactions we observed for SHIP1 were extremely transient in nature and inconsistent with the specificity and nanomolar affinities previously measured using surface plasma resonance (SPR), protein lipid overlay (PLO) assays, and nuclear magnetic resonance (NMR) spectroscopy (13–15).

We established conditions that enabled the detection of transient SHIP1 lipid interactions on supported membranes containing a mixture of 2% PI(3,4,5)P_3_ unimolecular kinetics and displayed no enhanced membrane recruitment over the course of any phosphatase catalyzed reaction. Overall, the lipid and 20% PS (τ_1_ = 25 ms, *K*_*OFF*_ = 40 sec^-1^). The dwell time was similar to those we measured for mEos-SHIP1 (PH-PP-C2) interacting with the plasma membrane in human neutrophils (**Figure 2I**). Since both PI(3,4,5) P_3_ and PS are required to detect a very transient interaction, the individual lipid interactions must have very weak affinities (*K*_*D*_ > 10 μM). This is consistent with our estimated *K*_*D*_ of 90 μM for the interaction between the SHIP1 active site and PI(3,4,5)P_3_, which we calculated based on a previously measured *K*_*M*_ (94 μM) and *k*_*cat*_ (8 sec^-1^) (32).

### SHIP1 activation by phosphatidylserine (PS)

The C2 domain of the SHIP1 paralog, SHIP2, reportedly interacts with phosphatidylserine (PS), which is proposed to allosterically stimulate phosphatase activity (16). Despite including up to 20% PS lipids in our TIRF-M supported membrane experiments, we did not observe an enhancement in the bulk membrane recruitment of mNG-SHIP1(PH-PP-C2). PS lipids did, however, stimulate the lipid phosphatase activity of SHIP1(PH-PP-C2), consistent with the reported mechanism of allosteric regulation for SHIP2 (16). Similarly, we observed a PS dependent increase in full-length SHIP1 activation. However, the overall activity was substantially lower compared to the SHIP1(PH-PP-C2) protein lacking autoinhibition. This suggests that full-length SHIP1 is autoinhibited by a mechanism that is partially resistant to PS-mediated activation. Considering that the plasma membrane contains 10-20% PS (33), full-length SHIP1 would be less susceptible to spuriously activation, which would help preserve cellular PI(3,4,5)P_3_ and PI(3,4)P_2_ lipid homeostasis. Based on our single molecule dwell time analysis of mNG-SHIP1(PH-PP-C2), the presence of both PS and PI(3,4,5)P_3_ lipids synergistically enhanced membrane binding. The transient interaction between the C2 domain and PS lipids could help orient the phosphatase domain on the membrane in a manner that makes SHIP1 more catalytically efficient. Looking at sequence homology between SHIP1 and SHIP2, we find that the phosphatase domains share 65% identity, while the C2 domains share 43% homology. Although PS-dependent allosteric activation of SHIP1 is potentially mediated by the phosphatase and C2 domain interface, like has been described for SHIP2 (16), the molecular basis remains unclear.

### Mechanism of SHIP1 autoinhibition

We performed a mutational analysis of full-length SHIP1 to determine which domains suppress lipid phosphatase activity. We found that deletion of the disordered C-terminus partially activates SHIP1, while deletion of the N-terminal SH2 domain fully activates the enzyme. Because the C-terminus of SHIP1 (892-1188aa) is predicted to be disordered, it remains unclear whether this region forms intramolecular contacts with a specific folded domain of SHIP1. Alternatively, the extended disordered C-terminal domain of SHIP1 could sterically hinder substrate binding. This type of mechanism has been reported for the guanine nucleotide exchange factor (GEF) Son of Sevenless (SOS), which similarly contains an extended disordered C-terminal domain and multiple layers of autoinhibition (34). Based on a biochemical assay used for measuring SOS dependent small GTPase activation, researchers previously reported that the presence of the unstructured poly-proline motif attenuated SOS GEF activity by lowering the association rate constant (*K*_*ON*_) of SOS (34). To date, no specific intramolecular interactions have been identified that regulate the C-terminal dependent autoinhibition of either SOS or SHIP1 activity.

Since the N- and C-terminus both contribute to autoinhibition, we initially hypothesized that SHIP1 was regulated by intramolecular interactions between the SH2 domain and tyrosine phosphorylated residues located in the C-terminus. By this mechanism, SHIP1 could adopt an autoinhibited confirmation similar to a Src-family non-receptor tyrosine kinase (20, 21). Suggestive of this model, the SH2 domain of SHIP1 was previously shown to interact with an isolated peptide containing phosphorylated Y1022 derived from SHIP1 (19). However, since these interactions were reconstituted using minimal fragments, it’s unclear whether the SH2 domain and the phospho-Y1022 peptide can form intramolecular contact in the context of length of SHIP1. Ultimately, determining the exact intramolecular interactions that regulate SHIP1 autoinhibition will require combining structural biochemistry and mutational analysis.

In our investigations of SHIP1 autoinhibition, we found that incubating SHIP1 with the promiscuous bacterial tyrosine phosphatase denoted YopH (35), did not relieve autoinhibition. This supports a model in which SHIP1 autoinhibition under our experimental conditions is not mediated by intramolecular interactions with phosphorylated tyrosine residues. However, in the context of immune cell signaling, post-translational modifications of the C-terminal NPxY motifs could cause SHIP1 to adopt an autoinhibited conformation that is mediated by intramolecular head-to-tail interactions or intermolecular daisy-chain structures.

In considering potential mechanisms for SHIP1 autoinhibition, we can draw parallels to the structural understanding of autoinhibited class 1A phosphatidylinositol 3-kinase (PI3K) (36). The regulatory subunit of PI3K contains two SH2 domains that each form intermolecular contacts with the catalytic subunit of PI3K. These interactions occur independently of a tyrosine phosphopeptide interaction. Similar to SHIP1, the SH2 domains of class I PI3Ks bind phosphorylated receptors, which releases autoinhibition to activate PI3K (36). Like PI3K, mutating residue R30A in SHIP1’s SH2 domain abolishes pY binding without disrupting autoinhibition. This supports that SHIP1 autoinhibition is mediated by the SH2 domain, but these interactions are distal to the pY binding pocket.

### Mechanisms controlling SHIP1 membrane localization in cells

The production of PI(3,4,5)P_3_ lipids during immune cell signaling has been shown to enhance plasma membrane localization of full-length SHIP1 (13). However, it has been unclear whether this response is regulated by a direct SHIP1-PI(3,4,5)P_3_ interaction or indirectly mediated by peripheral membrane binding proteins that bind to PI(3,4,5)P_3_ lipids. Single molecule visualization of SHIP1 in cells using photoconvertible Eos3.2 revealed that a small fraction of mEos-SHIP1(PH-PP-C2) molecules transiently localize to the plasma membrane. The membrane binding dynamics of mEos-SHIP1(PH-PP-C2) measured in cells were similarly transient to those measured on SLBs containing 2% PI(3,4,5)P_3_ and 20% DOPS. Consistent with membrane localization of purified mNG-SHIP1(PH-PP-C2) being insensitive to changes in PIP lipids composition *in vitro*, acute changes in PI(3,4,5)P_3_ levels did not modulate the bulk or single molecule membrane binding dynamics of SHIP1 in immune cells. Overall, the apparent lack of high affinity PS or PIP lipid interactions described here, suggests that SHIP1 membrane localization in cells is dominated by interactions with peripheral membrane proteins.

SHIP1 contains a variety of protein interaction domains and motifs that can potentially modulate its membrane localization and phosphatase activity in living cells. In the context of B-cell receptor signaling, SHIP1 is recruited by the ITIM motif of the inhibitory FcγRIIB receptor to down regulate this signaling process (31). SHIP1’s interaction with the FcγRIIB receptor is dependent on its SH2 domain and tyrosine phosphorylation of the ITIM motif (22). Our *in vitro* reconstitution demonstrates for the first time that membrane tethered phosphotyrosine peptides (i.e. pY-ITIM) can robustly recruit and activate SHIP1 on PI(3,4,5)P_3_ containing membranes. The data presented here also supports a model in which SHIP1 activity is largely regulated by its N-terminal SH2 domain. Although the C-terminus of SHIP1 has numerous protein interaction motifs that can potentially direct SHIP1 to critical locations at the plasma membrane, the molecular basis of those interactions is still being elucidated. By filling the gap in knowledge concerning how SHIP1 activity and membrane localization is regulated, in the future, we can better understand how cells control the strength and duration of PI(3,4,5)P_3_ signaling events. Given that SHIP1 is implicated in several diseases, including cancer (37–43), the described regulatory mechanisms will undoubtedly be of pharmacological interest.

## AUTHOR CONTRIBUTIONS

Resources: G.L.W., E.E.D., H.P.R., and S.D.H.

Experiments and investigation: G.L.W., E.E.D., H.P.R., and S.D.H.

Data Analysis: G.L.W., E.E.D., H.P.R., and S.D.H.

Conceptualization: G.L.W. and S.D.H.

Interpretation: G.L.W., E.E.D., H.P.R., and S.D.H.

Data curation: G.L.W. and S.D.H.

Writing – Review and editing: G.L.W., E.E.D., and S.D.H.

Writing – Original draft: G.L.W. and S.D.H.

Supervision: S.D.H.

Project administration: S.D.H.

Funding acquisition: S.D.H.

## MATERIALS & METHODS

### Molecular Biology

Gene coding for human phosphatidylinositol 4,5-bisphosphate phosphodiesterase delta-1 PH domain 11-140aa (PLCδ, Acc# P51178.2) was synthesized by GeneArt (Invitrogen) as a codon optimized open reading frame. The gene coding for the human tandem pleckstrin homology-domain-containing protein-1 PH domain (182-303aa, Uniprot #Q9HB21, *PKHA1*) was cloned using mKate2-P2A-APEX2-TAPP1-PH, a gift from Rob Parton (Addgene plasmid # 67662) (44). The gene coding for the Lactadherin-C2 phosphatidylserine lipid sensor, we cloned from Lact-C2-GFP, a gift from Sergio Grinstein (Addgene plasmid # 22852 ; http://n2t.net/addgene:22852; RRID:Addgene_22852) (45). The Lactadherin-C2 Domain was derived originally cloned from the *B. taurus* (bovine) gene called *MFGE8* (Uniprot #Q95114). Genes coding for human Src-homology-2-domain-containing inositol 5-phosphatase 1 (SHIP1 or INPP5D) was purchased as a cDNA clone from Dharmacon. Lentiviral packing vectors were purchased from Addgene. The psPAX2 plasmid was a gift from Didier Trono (Addgene plasmid #12260; http://n2t.net/addgene:12260; RRID:Addgene_12260). The pVSV-G plasmid was a gift from Akitsu Hotta (Addgene plasmid #138479; http://n2t.net/addgene:138479; RRID:Addgene_138479) (46). Gene fragments were subcloned into bacterial and baculovirus protein expression vectors containing coding sequences with different solubility and affinity tags. SHIP1 was cloned into a modified FAST Bac1 vector using Gibson Assembly. The complete open reading frame of all vectors used in this study were sequenced by Genewiz to ensure the plasmids lacked deleterious mutations. Each protein expression construct was screened for optimal yield and solubility in either bacteria (BL21 DE3 Star, Rosetta, etc.) or *Spodoptera frugiperda* (Sf9) insect cells.

### Purification of BACMID DNA

To create BACMID DNA, FASTBac1 plasmids were transformed into DH10 Bac cells and plated on agar containing 50 μg/mL kanamycin, 10 μg/mL tetracycline, 7 μg/mL gentamycin, 40 μg/mL X-GAL, and 40 μg/mL IPTG. After 2-3 days of growth at 37ºC, positive clones were isolated based on blue-white colony selection. Single white colonies were picked and restruck on a BACMID agar plate for a secondary selection. BACMIDs were purified from 3 mL bacterial cultures grown overnight in TPM. Bacteria are centrifuged and resuspended in 300 μL of buffer containing 50 mM Tris [pH 8.0], 10 mM EDTA, 100 μg/mL RNase A (Qiagen PI buffer). Bacteria were then lysed by adding 300 μL of buffer containing 200 mM NaOH, 1% SDS (Qiagen P2 buffer). Neutralize lysis buffer by adding 300 μL of 4.2 M Guanidine HCl, 0.9 M KOAc [pH 4.8] (Qiagen N3 buffer). Centrifuge sample at 23ºC for 10 minutes at 14,000 x g. Remove supernatant and combine with 700 μL 100% isopropanol. Centrifuge sample at 23ºC for 10 minutes at 14,000 x g. Remove supernatant and add 200 μL of 70% ethanol. Centrifuge sample at 23^0^C for 10 minutes at 14,000 x g. Remove supernatant and add 50 μL of 70% ethanol. Centrifuge sample at 23ºC for 10 minutes at 14,000 x g. Remove supernatant and dry DNA pellet slightly in biosafety hood. Solubilize DNA with 40 μL of sterile water. Resuspend DNA pellet by tapping side of micro-centrifuge tube 15-20 times. Quantify concentration of DNA using NanoDrop (typically 200-300 ng/μL). Immediately used BACMID DNA for transfection of Sf9 cells. Remaining BACMID DNA can be stored in -20ºC freezer.

### Baculovirus production

Baculoviruses was generated by transfecting 1 × 10^6^ *Spodoptera frugiperda* (Sf9) insect cells plated for 24 hours in a Corning 6-well plastic dish (Cat# 07-200-80) containing 2 mL of ESF 921 Serum-Free Insect Cell Culture media (Expression Systems, Cat# 96-001, Davis, CA.). All media contains 1x concentration of Antibiotic-Antimycotic (Gibco/Invitrogen, Cat#15240-062). For transfection, 5-7 μg BACMID DNA was incubated with 4 μL Fugene (ThermoFisher, Cat# 10362100) in 200 μL of ESF 921 media for 30 minutes at 23^0^C. BACMID DNA and Fugene were added dropwise to 6-well dish. Media change before and after addition of transfection reagent is unnecessary. After 4-5 days of transfection, viral supernatant (termed ‘P0’) is harvested, centrifuged, and used to infect 7 × 10^6^ *Sf9* cells plated for 24 hours in 10 cm tissue culture grade petri dish containing 10 mL of ESF 921 media and 10% Fetal Bovine serum (Seradigm, Cat# 1500-500, Lot# 176B14). After 4 days, viral supernatant (termed ‘P1’) is harvested and centrifuged to remove cell debri. Typical P1 viral titer yield is 10-12 mL. The P1 viral titer is expanded in 100 mL *Sf9* cell culture grown to a density of 1.25-1.5 × 10^6^ cells/mL in a sterile 250 mL polycarbonate Erlenmeyer flask with vented cap (Corning, #431144). We typically transduce 100 mL Sf9 culture with a concentration of 1% vol/vol of PI viral titer. Remaining PI virus is frozen as 1.5 mL aliquots that are stored in the -80^0^C freezer. The 10% Fetal Bovine serum serves as a cryo-protectant. After 4 days, viral supernatant (termed ‘P2’) is harvested, centrifuged, and 0.22 μm filtered in 150 mL filter-top bottle (Corning, polyethersulfone (PES), Cat#431153). The P2 viral supernatant is used for protein expression in either *Sf9* or High 5 cells grown in ESF 921 Serum-Free insect cell culture media. The MOI for optimal protein expression is determined empirically to minimize cell death and maximize protein yield (typically 1.5-2% vol/vol final concentration of P2 virus).

### Purification of SHIP1 and SHIP1 mutants (FL, ΔCT, ΔSH2)

The coding sequence of full-length human SHIP1 (1-1188aa) was cloned into a modified FastBac1 vector containing an N-terminal his6-TEV-mNG fusion. High five insect cells were infected with baculovirus using an optimized multiplicity of infection (MOI), typically 1.5–2% vol/vol, which we empirically determined based on small-scale test expressions (25-50 mL culture). Infected cells were typically grown for 48 hours at 27ºC in ESF 921 Serum-Free Insect Cell Culture medium (Expression Systems, Cat# 96-001-01). Cells were harvested by centrifugation, transferred to 50 mL tubes by washing with 1x PBS [pH 7.2], pelleted and resuspended in 10 mL of 1x PBS [pH 7.2], 10% Glycerol, and 2x protease inhibitor cocktail (1 Sigma protease inhibitor tablet per 50 mL of buffer) and then stored in the -80ºC freezer. For purification, frozen cell pellets were thawed in an ambient water bath and lysed into buffer containing 30 mM Tris [pH 8.0], 10 mM imidazole, 400 mM NaCl, 1 mM PMSF, 2 mM BME, Sigma protease inhibitor cocktail EDTA free per 100 mL lysis buffer, and 100 μg/mL DNase using a dounce homogenizer. Lysate was then centrifuged at 37,000 rpm (140,000xg) for 60 minutes in a Beckman Ti70 rotor at 4ºC. High speed supernatant (HSS) was then batch bound to 5 mL of Ni-NTA Agarose (Qiagen, Cat# 30230) resin at 4ºC for 2 hours stirring in a beaker. The resin and HSS was collected in 50 mL tubes, centrifuged, and washed with buffer containing 20 mM Tris [pH 8.0], 30 mM imidazole, 300-400 mM NaCl, 5% Glycerol, and 2 mM BME before being transferred to gravity flow column. NiNTA resin with bound his6-TEV-mNG-SHIP1 was then washed with an additional 100 mL of 20 mM Tris [pH 8.0], 30 mM imidazole, 300-400 mM NaCl, 5% Glycerol, and 2 mM BME buffer and then eluted into buffer containing 300 mM imidazole. Peak fractions were pooled and desalted using a G25 Sephadex desalting column (GE Healthcare/Cytiva) equilibrated in 20 mM Tris [pH 8.0], 100 mM NaCl, and 1mM TCEP buffer. Any precipitation was removed via 0.22 μm syringe filtration. Clarified SHIP1 fractions were then bound to a MonoQ anion exchange column (GE Healthcare/Cytiva) equilibrated in 20 mM Tris [pH 8.0], 100 mM NaCl, and 1 mM TCEP buffer. Proteins were resolved over a 10-100% linear gradient (0.1-1 M NaCl, 30 minutes, 1.5 mL/min flow rate). His6-TEV-mNG-SHIP1 elutes in the presence of 200-400 mM NaCl. Peak fractions containing SHIP1 were pooled, incubated with 400 μL of 243 μM his6-TEV protease (S291V) with poly-R tail for ∼12-16 hours at 4^0^C, concentrated in a 30 kDa MWCO Amicon concentrator (Sigma Millipore), and then loaded onto a 24 mL Superdex 200 10/300 GL (GE Healthcare, Cat# 17-5174-01) size exclusion column equilibrated in 20 mM Tris pH [8] (20mM HEPES pH [7.5] for FL SHIP1), 150 mM NaCl, 10% glycerol, 1 mM TCEP, and 0.5% CHAPS buffer. Peak fractions were concentrated in a 30 kDa MWCO Vivaspin 6 centrifuge tube and flash frozen at a final concentration of 1-30 μM using liquid nitrogen. Purification schemes for ΔSH2, ΔCT, and FL SHIP1 were nearly identical. ΔSH2 SHIP1 was incubated with 400 μL of 243 μM his6-TEV protease (S291V) with poly-R tail for ∼12-16 hours at 4ºC before Desalting, MonoQ, and S200 chromatography steps.

### SHIP1 mutant (mNG-PH-PP-C2)

BL21 (DE3) Star bacteria were transformed with his10-TEV-mNG-GGGGG-SHIP1 (mNG-PH-PP-C2, 191-878aa) and plated on LB agar containing 50 μg/ml kanamycin. The following day, 50mL of TPM media (20g tryptone, 15g yeast extract, 8g NaCl, 2g Na_2_HPO_4_, and 1g KH_2_PO_4_ per liter) containing 50 μg/ml kanamycin was inoculated with one bacterial colony from the agar plate. This culture was grown for 12-16 hours at 27°C to an OD600 = 2–3, before being diluted to an OD600 = 0.05–0.1 in 2 liters of TPM media. These cultures were grown at 37°C to an OD600 = 0.6-0.8, shifted to 30°C for 1 hour, and then bacteria were induced with 50 μM IPTG to express SHIP1 mutants. After 6-8 hours of expression at 30°C, bacterial cultures were harvested by centrifugation, transferred to 50 mL tubes by washing with 1x PBS [pH 7.2], pelleted, and then stored in the -80°C freezer.For purification, frozen cell pellets were thawed in an ambient water bath and lysed into buffer containing 50 mM Na_2_HPO_4_ [pH 8.0], 400 mM NaCl, 1 mM PMSF, 0.4 mM BME, and 100 μg/mL DNase by micro-tip sonication (12-24 × 5 second pulses, 40% amplitude). Lysate was then centrifuged at 15,000 rpm (35,000xg) for 60 minutes in a Beckman JA-20 rotor at 4ºC. During lysate centrifugation, a 5mL HiTrap chelating column (Cat#) was charged with 50 mL of 100mM CoCl_2_, generously washed with water (phosphate buffer will precipitate unchelated CoCl_2_ if not thoroughly removed) and equilibrated with lysis buffer lacking PMSF and DNase (>0.4mM BME will destroy charged HiTrap column and turn column brown or black). High speed supernatant (HSS) was then circulated over charged 5mL HiTrap column for 2 hours. Captured protein was washed with 100mL of 50 mM Na_2_HPO_4_ [pH 8.0], 400 mM NaCl, 10mM imidazole, and 0.4 mM BME and then eluted into buffer containing 500 mM imidazole. Peak fractions containing SHIP1 were pooled with 400 μL of 243 μM his6-TEV protease (S291V) with poly-R tail and dialyzed in 4L of 25 mM Na_2_HPO_4_ [pH 8.0], 400 mM NaCl, and 0.4 mM BME buffer at 4°C for ∼12-16 hours to remove imidazole. The next day, the charged HiTrap column was equilibrated with dialysis buffer and dialyzed protein was recirculated for 2 hours at 4°C to remove his6-TEV protease and any uncleaved his10-TEV-mNG-GGGGG-SHIP1. Unbound protein was then concentrated in a 30 kDa MWCO Amicon concentrator (Sigma Millipore) and then loaded onto a 24 mL Superdex 200 10/300 GL (GE Healthcare, Cat# 17-5174-01) size exclusion column equilibrated in 20 mM Tris pH [8], 200 mM NaCl, 10% glycerol, and 1 mM TCEP buffer. Peak fractions were concentrated in a 30 kDa MWCO Vivaspin 6 centrifuge tube and flash frozen at a final concentration of 10-30 μM using liquid nitrogen.

### Sortase mediated peptide ligation

All lipid sensors were labeled on a N-terminal (Gly)_5_ motif using sortase mediated peptide ligation (Ton-That et al. 1999; Guimaraes et al. 2013). Hansen et al. devised a novel approach for chemically modifying an LPETGG peptide with fluorescent dyes, which we then conjugated to our protein of interest. The LPETGG peptide was synthesized to >95% purity by ELIM Biopharmaceutical (Hayward, CA) and labeled on the N-terminal amine with N-Hydroxysuccinimide (NHS) fluorescent dye derivatives (e.g. NHS-Alexa488). This was achieved by combining 10 mM LPETGG peptide, 15 mM NHS-Alexa488 (or other fluorescent derivatives), and 30 mM Triethylamine (Sigma, Cat# 471283) in anhydrous DMSO (Sigma, Cat# 276855). This reaction was incubated overnight in the dark at 23ºC before being stored in a -20ºC freezer. Prior to labeling (Gly)_5_ containing proteins, unreacted NHS-Alexa488 remaining in the LPETGG labeling reaction was quenched with 50 mM *Tris*(hydroxymethyl) aminomethane (Tris) [pH 8.0] buffer for at least 6 hours. Complete quenching of unreacted NHS-Alexa488 was verified by the inability to label (Gly)_5_ containing proteins in the absence of a Sortase. When labeling (Gly)_5_ containing proteins with the fluorescently labeled LPETGG peptide we typically combined the following reagents: 50 mM Tris [pH 8.0], 150 mM NaCl, 50 μM (Gly)_5_-protein, 500 μM Alexa488-LPETGG, and 10-15 μM His_6_-Sortase (Δ1-57aa). This reaction mixture was incubated at 16-18ºC for 16-20 hours, before buffer exchange with a G25 Sephadex column (e.g. PD10 or NAP5) to remove majority of dye and dye-peptide. The his_6_-Sortase was then captured on NiNTA agarose resin (Qiagen) and unbound, labeled protein was separated from remaining fluorescent dye and peptide using a Superdex 75 or Superdex 200 size exclusion chromatography column (24 mL bed volume).

### Preparation of small unilamellar vesicles

The following lipids were used to generated small unilamellar vesicles (SUVs): 1,2-dioleoyl-sn-glycero-3-phosphocholine (18:1 DOPC, Avanti # 850375C), 1,2-dioleoyl-*sn-*glycero-3-phospho-L-serine (18:1 DOPS, Avanti # 840035C), 1,2-dioleoyl-sn-glycero-3-phosphoethanolamine-N-[4-(p-maleimidomethyl)cyclohexane-carboxamide] (18:1 MCC-PE, Avanti # 780201C), D-myo-phosphatidylinositol 3,4,5-trisphosphate (PI(3,4,5)P_3_ diC16, Echelon P-3916-100ug). Lipids from Avanti were purchased as single use ampules containing between 0.1-5 mg of lipids dissolved in chloroform. PI(3,4,5)P_3_ lipids from Echelon were purchased as salts and solubilized in the manufacturer’s suggested solvent containing 263 μL CHCl_3_, 526 μL MeOH, and 211 μL H_2_O per mL volume. To make liposomes, 2 μmoles total lipids are combined in a 35 mL glass round bottom flask containing 2 mL of chloroform. Lipids are dried to a thin film using rotary evaporation with the glass round-bottom flask submerged in a 42ºC water bath. After evaporating all the chloroform, the round bottom flask was place under a vacuum for 15 minutes. The lipid film was then resuspended in 2 mL of PBS [pH 7.4], making a final concentration of 1 mM total lipids. All lipid mixtures expressed as percentages (e.g. 98% DOPC, 2% PI(3,4,5)P_3_) are equivalent to molar fractions. For example, a 1 mM lipid mixture containing 98% DOPC and 2% PI(3,4,5)P_3_ is equivalent to 0.98 mM DOPC and 0.02 mM PI(3,4,5)P_3_. To generate 30-50 nm SUVs, 1 mM total lipid mixtures were extruded through a 0.05 μm pore size 19 mm polycarbonate membrane (Sigma, cat#WHA800308) with filter supports (Avanti, cat#610014) on both sides of the PC membrane. Hydrated lipids at a concentration of 1 mM were extruded through the PC membrane 11 times.

### Preparation of supported lipid bilayers

Supported lipid bilayers are formed on 25×75 mm coverglass (IBIDI, #10812). Coverglass was first cleaned with 2% Hellmanex III (ThermoFisher, Cat#14-385-864) heated to 60-70ºC in a glass coplin jar for at least 30 minutes. Coverglass is then washed extensively with MilliQ water and then etched with Pirahna solution (1:3, hydrogen peroxide:sulfuric acid) for 5-10 minutes the same day SLBs were formed. Etched coverglass, washed extensively with MilliQ water again, is rapidly dried with nitrogen gas before adhering to a 6-well sticky-side chamber (IBIDI, Cat# 80608). SLBs are formed by flowing 30 nm SUVs diluted in 1x PBS pH [7.2] to a total lipid concentration of 0.25 mM into the assembled chamber. After 30 minutes, IBIDI chambers are washed with 4 mL of PBS [pH 7.4] to remove non-absorbed SUVs. Membrane defects are blocked for 10 minutes with a 1 mg/mL beta casein (ThermoFisher, Cat# 37528) diluted in 1x PBS [pH 7.4]. Before it was used as a blocking protein, frozen 10 mg/mL beta casein stocks were thawed, centrifuged for 30 minutes at 21370 x *g*, and 0.22 μm syringe filtered. After blocking SLBs with beta casein, membranes were washed again with 1mL of 1x PBS, followed by 1 mL of kinase buffer (20 mM HEPES [pH 7.0], 150 mM NaCl, 1 mM ATP, 5 mM MgCl_2_, 0.5 mM EGTA, 200 μg/ mL beta casein, 20 mM BME, and 20 mM glucose) before TIRF-M.

### Conjugation of pY peptide to supported membranes

SLBs with MCC-PE lipids were used to covalently couple phosphorylated peptides to the maleimide lipid headgroups. Phosphorylated peptides (**C**KTEAENTIT(pY)SLIK, Elim Biopharmaceuticals) correspond to the ITIM sequence of the FcγRIIB C-terminus (19). A cysteine (bold) was added to the N-terminus of the peptide to form a covalent interaction with the maleimide lipid headgroup of the MCC-PE lipids. We refer to this peptide as pY-ITIM throughout the manuscript. For these SLBs, 100 μL of 10 μM pY-ITIM peptides diluted in a 1x PBS [pH 7.4] and 0.1 mM TCEP buffer was added to the IBIDI chamber and incubated for 2 hours. The addition of TCEP significantly increases the coupling efficiency. SLBs with MCC-PE lipids were then washed with 2 mL of 5mM BME diluted in 1x PBS [pH 7.4] and incubated for 15 minutes to destroy the unreacted maleimide headgroups. SLBs were washed with 1mL of 1x PBS, followed by 1 mL of kinase buffer before TIRF-M.

### Oxygen scavenging system

Glucose oxidase (32 mg/mL, 100x stock) and catalase (5 mg/mL, 100x stock) were solubilized in 20 mM HEPES [pH 7.0], 150 mM NaCl, 10% glycerol, and 1 mM TCEP buffer, and then flash frozen in liquid nitrogen and stored at -80°C. Trolox (200mM, 100x stock) is UV treated and stored at -20°C. Approximately 10 minutes before imaging, 100x oxygen scavenger stocks were diluted in kinase buffer containing enzymes/biosensors to achieve a final concentration of 320 μg/mL glucose oxidase (Serva, #22780.01 *Aspergillus niger*), 50 μg/mL catalase (Sigma, #C40-100MG Bovine Liver), and 2 mM Trolox (Cayman Chemicals, cat #10011659). Trolox was prepared using a previously described protocol that utilized UV irradiation to drive the formation of a quinone species (47, 48).

### Kinetics measurements of PI(3,4)P_2_ production

The kinetics of PI(3,4,5)P_3_ dephosphorylation via SHIP1 was measured on SLBs formed in IBIDI chambers and visualized using TIRF microscopy. The reaction buffer contains 20 mM HEPES [pH 7.0], 150 mM NaCl, 1 mM ATP, 5 mM MgCl_2_, 0.5 mM EGTA, 200 μg/mL beta casein, 20 mM BME, and 20 mM glucose. Most kinetics assays used initial membrane compositions of 96% DOPC, 2% PI(3,4,5)P_3_, and 2% MCC-PE, or 86% DOPC, 10% DOPS, 2% PI(3,4,5)P_3_, 2% MCC-PE. Membrane compositions that varied from the above conditions are described in figure legends. As indicated in the figure legend, we measured the conversion PI(3,4,5)P_3_ to PI(3,4) P_2_ by visualizing the membrane localization of either 20 nM Alexa647-Grp1 (PH domain; PI(3,4,5)P_3_ sensor) or 6 nM Cy3-TAPP1 (PH domain; PI(3,4)P_2_ sensor) with TIRF-M. By convention, enzyme kinetics are plotted in terms of product formation rather than substrate depletion. For experiments that utilize the Grp1 sensor, normalized kinetic traces were inverted to reflect PI(3,4)P_2_ production. Assuming a footprint of 0.72 nm^2^ as reported for DOPC lipids (49), we calculated a density of 27778 lipids/μm^2^ for 2% PI(3,4)P_2_. Apparent rate constants (k_app_) were calculated using the half-time relationship for first-order reactions: t_1/2_ = 0.693/k_app_.

### Cell culture

PLB-985 neutrophil-like cells are an established model cell line for human neutrophils and were obtained from the laboratory of Dr. Sean Collins (University of California at Davis). PLB-985 cells were grown in suspension in RPMI 1640 + GlutaMAX media containing 25 mM HEPES (Life Technologies, cat #72400047), 9% fetal bovine serum (FBS), penicillin (100 units/ml) (Life Technologies, cat #15140122), streptomycin (100 μg/ml) (Life Technologies, cat #15140122). Cell lines were grown in humidified incubators at 37°C in the presence of 5% CO2 and split three times per week, keeping densities between 0.1-2 × 10^6^ cells/mL. PLB-985 cells were differentiated into a neutrophil-like state by culturing 0.2 × 10^6^ cells/mL for 6 to 7 days in RPMI media supplemented with 2% FBS, penicillin (100 units/ml), streptomycin (100 μg/ml), 1.3% DMSO, and 2% Nutridoma-CS (Sigma, cat #11363743001). Nutridoma-CS was added to increase the chemotactic response of the cells, but this supplement is optional (50). Human embryonic kidney (HEK) 293T Lenti-X were obtained from Takara (cat# 632180). These cells transformed with adenovirus type 5 DNA and expresses the SV40 large T antigen. HEK293T Lenti-X cells were cultured in DMEM + GlutMAX + High Glucose (4.5 g/L) + sodium pyruvate (110 mg/L) (Life Technologies, cat #10569010) supplemented with 10% FBS (Sigma, cat# F4135-500ML), penicillin (100 units/ml), and streptomycin (100 μg/ml). Cells were grown in 10 cm dishes in humidified incubators at 37°C in the presence of 5% CO_2_ and split at a confluency of 80-90% every 2-3 days. HEK293T Lenti-X cells were split using using 1.5mL of 0.25% Trypsin. Trypsin was that quenched with 8.5 mL complete DMEM media containing 10% FBS. Cells were diluted 1:10 and seeded on a new 10cm dish containing a total volume of 10 mL complete DMEM media warmed to 37°C.

### Lentivirus production

Lentivirus was generated by transfecting 60-70% confluent HEK293 Lenti-X cells in a 10-cm plate containing 8 mL of complete media. Transfection reagents were prepared by combining 6.7 μg psPAX2 (2nd generation lentiviral packaging plasmid, Addgene #12260), 0.85 μg pVSV-G (Expresses VSV-G envelop protein for pseudotyping NanoMEDIC particle, Addgene #138479) (46), 7.5 μg of transfer lentiviral vector containing gene of interest, and 30 μL of 1 mg/mL polyethyleneimine (PEI) in 0.5 mL Opti-Mem (ThermoFisher, cat#31985070). This mixture was incubated for 15 minutes at room temperature before adding dropwise to plated HEK293 Lenti-X cells. Media containing lentivirus was harvested 48 hours after initiating transfection and clarified by centrifugation to remove cell debris. Lentivirus was concentrated by adding 333 μL Lenti-X concentrator (Takara, cat# 631231) per mL of viral supernatant, incubated overnight at 4°C, and then centrifuged at 1500 x g for 45 minutes. Supernatant was aspirated and the white viral containing pellet was resuspended in 0.4 mL of complete RPMI media (9% FBS). Lentivirus was used immediately or stored at -80°C. To infect PLB-985 cells, 0.4 mL of concentrated lentivirus was added with 5 mL of undifferentiated cells at a density of 0.2 × 10^6^ cells/mL containing a final concentration of 8 μg/mL polybrene (Millipore, Cat# TR-1003-G, 10 mg/mL stock, 1250x) in the cell culture media. The cells were passaged at least one time before differentiating into the neutrophil-like state (see ***Cell Culture*** for protocol).

### Live cell imaging

Extracellular Buffer (ECB: 5mM KCl, 125 mM NaCl, 1.5 mM CaCl2, 1.5 mM MgCl2, 10 mM glucose, 20 mM HEPES pH 7.4) and Leibovitz-15 (Life Technologies, cat #21083027) complete media (10% FBS, 100 U/mL Pen/ Strep) were warmed to 37°C and combined 1:4 to make the Imaging Media. Differentiated PLB-985 cells were prepared for imaging by centrifuging 0.5 mL of cells at 100 x *g* for 10 minutes, aspirating off the medium, and resuspending the cells in warm imaging media (1:4 mixture of L-15 complete media and ECBaq). Differentiated PLB-985 cells were then imaged using one of two different live cell imaging methods: a method using fibronectin coated glass attached to an IBIDI flow cell chamber or a previously described 96-well format under-agarose method (51). To image cells, 25 × 75 mm coverslips were cleaned with 2% Hellmanex III (ThermoFisher, Cat#14-385-864), washed with MilliQ water, dried with N_2_ gas, and attached to an IBIDI flow cell chamber (IBIDI sticky-Slide VI 0.4, cat #80608). A 10 μg/mL fibronectin (Sigma, cat#F1141, 1 mg/mL stock concentration) solution diluted in 1x PBS was added to each well of the IBIDI chamber and incubated for 60 minutes at 23ºC. Unbound fibronectin was washed out with 1x PBS and then blocked with 0.2% BSA in 1x PBS (bovine serum albumin fatty/endotoxin free, Sigma, cat#) for 15 minutes. Differentiated PLB-985 cells centrifuged and suspended in imaging media (recipe above) were flowed into the IBIDI chamber and allowed to adhere for 10-15 minutes. Cells were visualized by TIRF-M in imaging media and stimulated with a final uniform concentration of 20 nM fMLF (N-Formyl-L-methionyl-L-leucyl-L-phenylalanine, Sigma, cat#F3506) to drive PI(3,4,5)P_3_ production.

### Microscope hardware and imaging acquisition

Membrane binding and lipid phosphorylation reactions reconstituted on supported lipid bilayers (SLBs) were visualized using an inverted Nikon Eclipse Ti2 microscope using a 100x Nikon (1.49 NA) oil immersion TIRF objective. TIRF-M images of SLBs were acquired using an iXion Life 897 EMCCD camera (Andor Technology Ltd., UK). Fluorescently labeled proteins were excited with either a 488 nm, 561 nm, or 637 nm diode laser (OBIS laser diode, Coherent Inc. Santa Clara, CA) controlled with a Vortran laser drive with acousto-optic tunable filters (AOTF) control. The power output measured through the objective for single particle imaging was 1-2 mW. Excitation light was passed through the following dichroic filter cubes before illuminating the sample: (1) ZT488/647rpc and (2) ZT561rdc (ET575LP) (Semrock). Fluorescence emission was detected on the iXion Life 897 EMCCD camera position after a Nikon emission filter wheel housing the following emission filters: ET525/50M, ET600/50M, ET700/75M (Semrock). All experiments were performed at room temperature (23ºC) for both biochemistry and live-cell imaging. Microscope hardware was controlled by Nikon NIS elements.

### Single particle tracking algorithm

After the 16-bit .tif images were cropped down to 400×400 to minimize differences in field illumination. Upon starting the TrackMate plugin (52) on ImageJ/Fiji, the initial calibration settings were default and enter ‘YES’ for allow swapping of Z or T (depth or time). Select the LoG detector and enter the appropriate protein-related dimensions. Check the boxes applying median filter and sub-pixel localization. Apply default settings for initial thresholding, run hyperstack display, and then filter particles based on mean intensity. Choose a tracker (simple LAP tracker was used in this paper, but LAP tracker also works). Set the following filters on the tracks: Track Start (remove particles at start of movie); Track End (remove particles at end of movie); Track displacement; X - Y location (above and below totaling 4 filters). Continue running program and display options according to one’s preference. From the generated files, the dwell times and step sizes were extracted manually. A histogram and then a cumulative distribution was generated for each data set. Dwell times were fitted with a one or two species exponential function.

To calculate the dwell times for membrane bound molecules, we sorted into a cumulative distribution frequency (CDF) plot with the frame interval as the bin (e.g. 8-50 ms). The log_10_(1-CDF) is then plotted against the dwell time and fit to a single or double exponential.

Single exponential model:

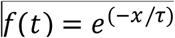

Two exponential model:

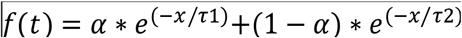

Fitting procedure initiated with a single exponential. In cases of a low-quality single exponential fit, a maximum of two populations were used. For double exponential fit, alpha represents the percentage of the fast-dissociating molecules characterized by τ_1_.

