## Supplemental Figures for "Mechanisms controlling membrane recruitment and activation of autoinhibited SHIP1"

### Supplemental Figure 1

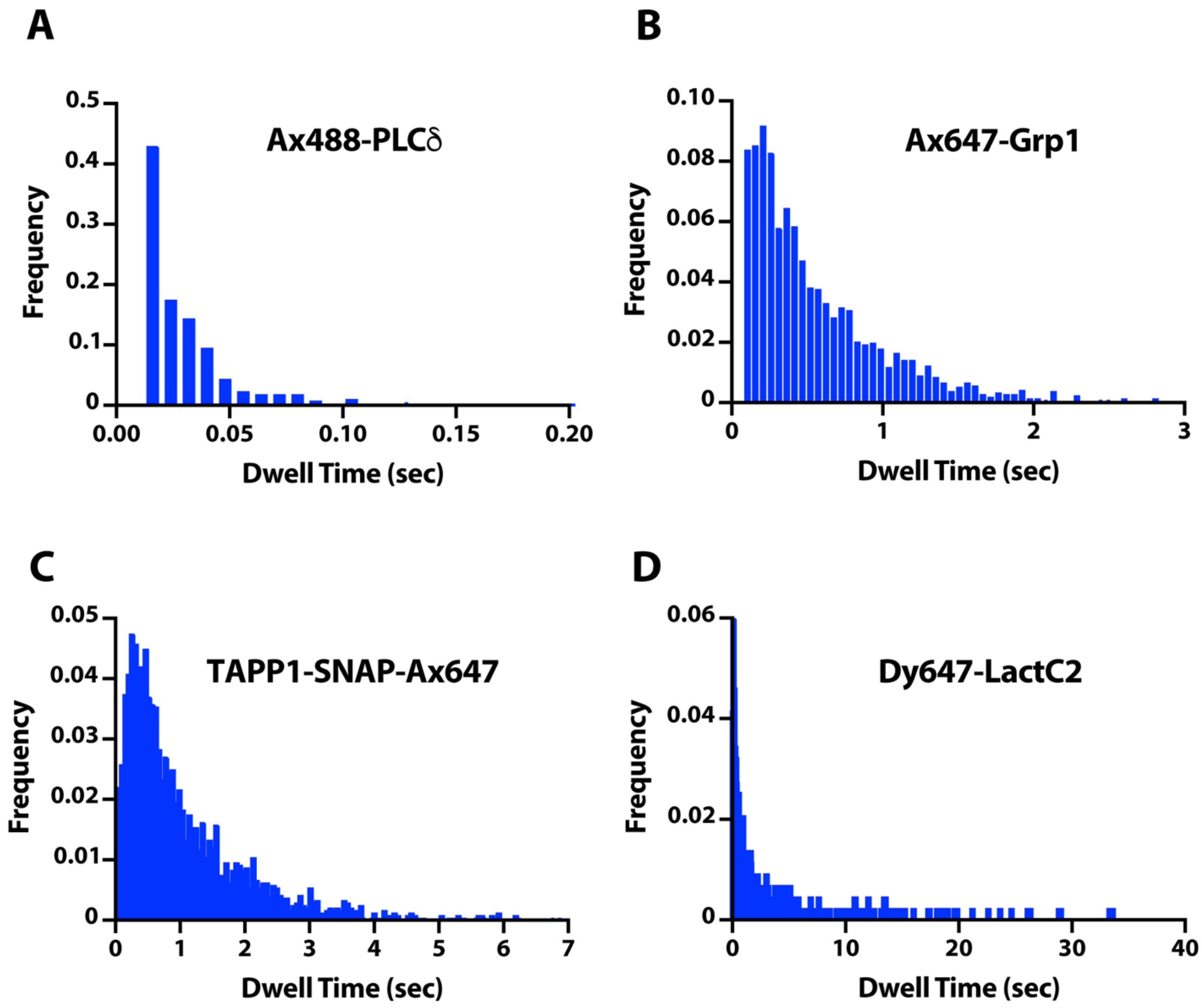

**Supplementary Figure 1**
**Single molecule dwell time distributions of PIP lipid binding domains measured on supported membranes using smTIRF-M**
**(A-D)** Representative dwell time frequency distributions for data shown in Figure 1E. Data was collected under the following conditions: **(A)** 250 pM Ax488-PLCδ + 2% PI(4,5)$P_2$, **(B)** 1 pM Ax647-Grp1 + 2% PI(3,4,5)$P_3$, **(C)** 1 pM TAPP1-SNAP-Ax647 + 2% PI(3,4)$P_2$, **(D)** 2 pM LactC2-Dy647 + 20% DOPS. This data is presented in Figure 1E yielding the following dwell times: Ax488-PLCδ ($\tau_1$ = 24 ± 2 ms) TAPP1-SNAP-Ax647 ($\tau_1$ = 1.02 ± 0.053 s), Ax647-Grp1 ($\tau_1$ = 0.544 ± 0.007 s), or LactC2-Dy647 ($\tau_1$ = 0.765 ± 0.191 s, $\tau_2$ = 6.58 ± 0.539 s, α = 0.5 ± 0.04). Note that the exponential curve fits are not shown. Membrane composition: DOPC lipids plus the indicated PIP lipids concentration stated above.

### Supplemental Figure 2

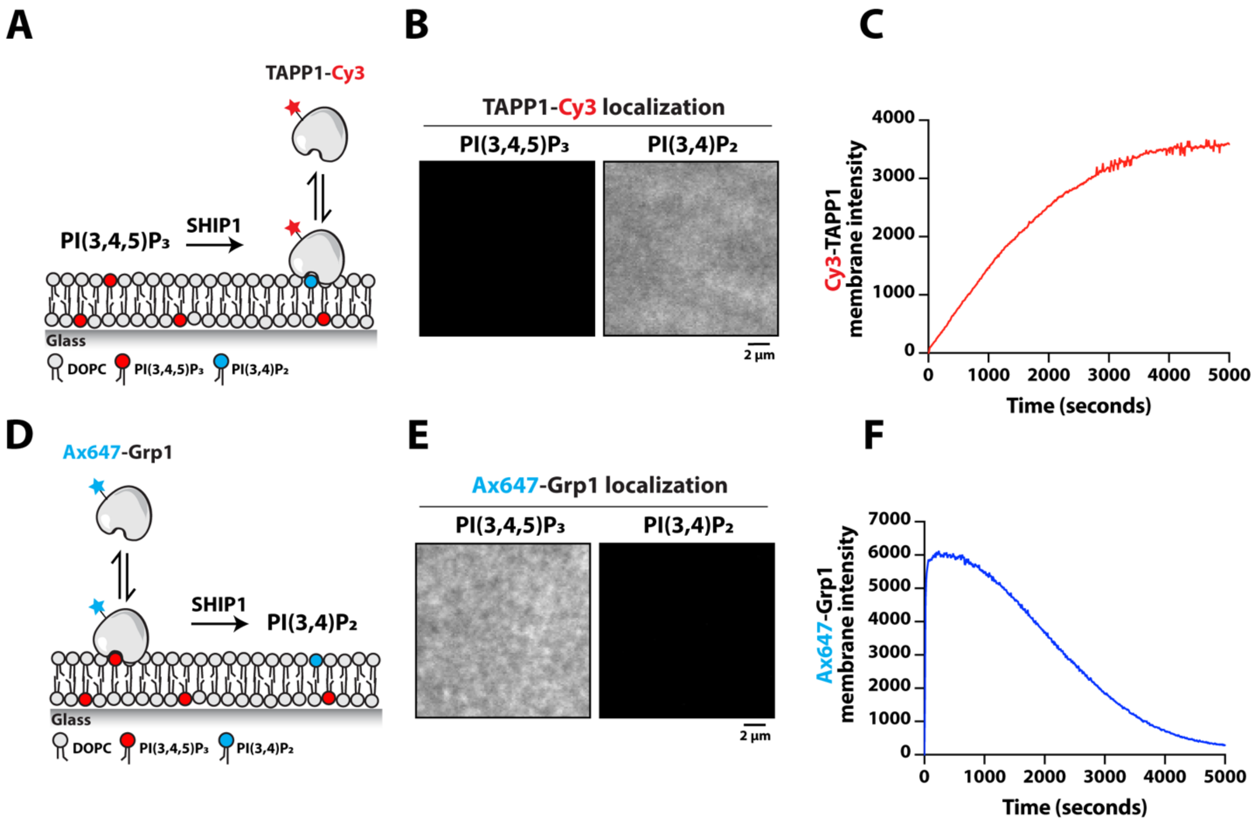

**Supplementary Figure 2**

**Validation of TAPP1 and Grp1 sensors for measuring dephosphorylation of PI(3,4,5)$P_3$**

**(A)** Cartoon schematic showing membrane association of Cy3-TAPP1 in a PI(3,4)$P_2$ dependent manner. **(B)** Representative TIRF-M images showing the localization of 20 nM Cy3-TAPP1 on SLBs containing 98% DOPC with either 2% PI(3,4,5)$P_3$ or 2% PI(3,4)$P_2$. **(C)** Kinetics of PI(3,4,5)$P_3$ dephosphorylation measured in the presence of 50 nM full-length SHIP1 and 50 nM Cy3-TAPP1. **(D)** Cartoon schematic showing membrane association of Alexa647-Grp1 in a PI(3,4,5)$P_3$ dependent manner. **(E)** Representative TIRF-M images showing the localization of 20 nM Alexa647-Grp1 on SLBs containing 98% DOPC with either 2% PI(3,4,5)$P_3$ or 2% PI(3,4)$P_2$. **(F)** Kinetics of PI(3,4,5)$P_3$ dephosphorylation measured in the presence of 50 nM full-length SHIP1 and 20 nM Alexa647-Grp1.

### Supplemental Figure 3

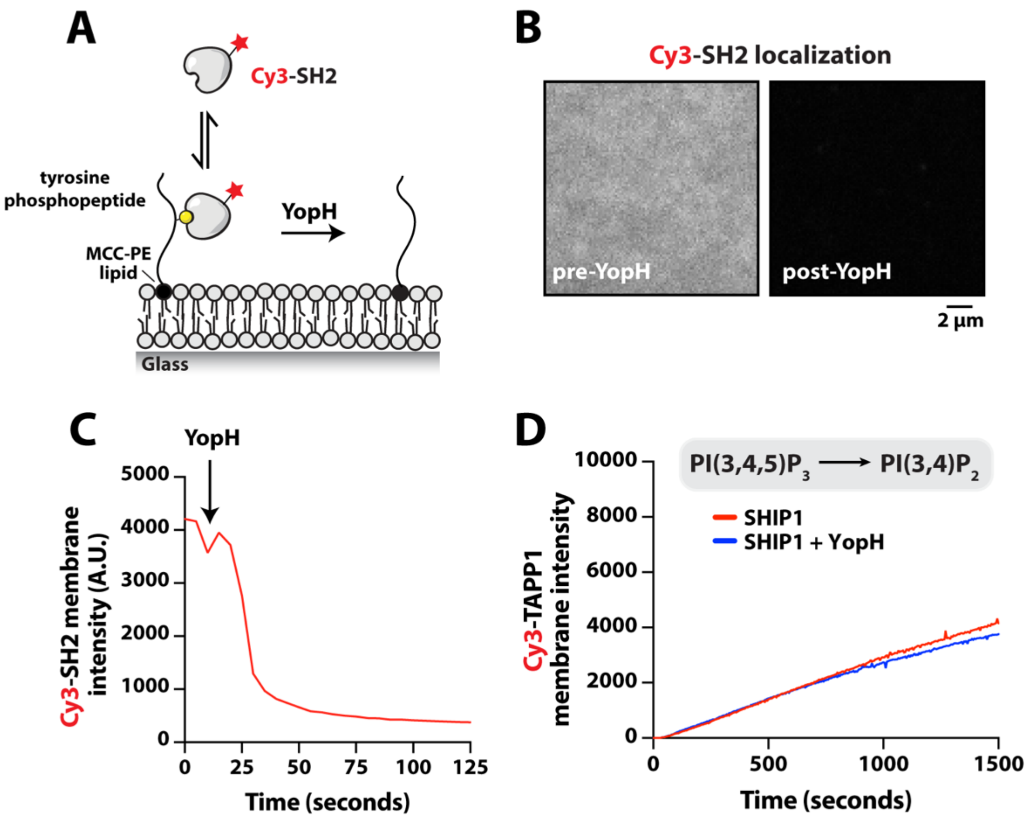

**Supplementary Figure 3**
**Tyrosine phosphatase, YopH, does not stimulate activity of full-length SHIP1**
**(A)** Cartoon schematic showing membrane association of Cy3-SH2 with membrane anchored phosphotyrosine peptide derived from a ITIM motif (pY-ITIM). The tyrosine phosphatase, YopH, drives the dissociation of Cy3-SH2 following dephosphorylation of the pY-ITIM peptide. **(B)** Representative TIRF-M images showing the localization of 50 nM Cy3-SH2 in the absence and presence of 10 nM YopH. **(C)** Kinetics of phosphotyrosine (pY-ITIM) peptide dephosphorylation monitored in the presence of 50 nM Cy3-SH2 and 10 nM YopH. **(D)** Full-length SHIP1 is not activated by the addition of the tyrosine phosphatase YopH. Reactions contained 50 nM SHIP1 (FL), 20 nM Cy3-TAPP1, 50 nM YopH was incubated for 10 minutes before injecting in SLB chamber and imaged using TIRF-M. **(B-D)** Membrane composition: 96% DOPC, 2% PI(3,4,5)$P_3$, 2% MCC-PE-(pY conjugated).

### TABLE S1

| Protein visualized | Membrane composition | $\tau_1$ ± *SD (sec)* | $\tau_2$ ± *SD (sec)* | $\alpha$ ± *SD* | *N* | *n* |
|---|---|---|---|---|---|---|
| Alexa488-PLC$\delta$ | 2% PI(4,5)$P_2$ | 0.024 ± 0.002 | — | — | 4 | 389 |
| Alexa647-Grp1 | 2% PI(3,4,5)$P_3$ | 0.544 ± 0.005 | — | — | 3 | 5588 |
| TAPP1-SNAP-Alexa647 | 2% PI(3,4)$P_2$ | 1.02 ± 0.053 | — | — | 3 | 8102 |
| LactC2-Dy647 | 20% PS | 0.765 ± 0.191 | 6.58 ± 0.539 | 0.50 ± 0.04 | 3 | 1167 |
| mEos-LactC2 | PM | 0.371 ± 0.069 | — | — | 12 | 18124 |
| mEos-Grp1 | PM | 0.392 ± 0.048 | — | — | 10 | 10184 |
| mEos-SHIP1 (PH-PP-C2) | PM | 0.038 ± 0.003 | — | — | 4 | 8480 |
| mEos-SHIP1 (PH-PP-C2) | PM + fMLF | 0.037 ± 0.005 | — | — | 5 | 7975 |
| mNG-SHIP1 (PH-PP-C2) | 2% PI(3,4,5)$P_3$, 20% PS | 0.025 ± 0.001 | — | — | 2 | 1562 |
| mNG-SHIP1 (PH-PP-C2) | 2% PI(3,4,5)$P_3$, 20% PS* | 0.009 ± 0.002 | 0.056 ± 0.007 | 0.44 ± 0.13 | 5 | 4801 |

SD = standard deviation from the indicated number of technical replicates
*N* = # of SLBs or cells used for calculating the mean dwell times (i.e. technical replicates)
n = total number of molecules tracked across the indicated number of technical replicates (*N*)
alpha ($\alpha$) = fraction of molecules with characteristic dwell time ($\tau_1$ and $\tau_2$)
membrane composition: DOPC plus the indicated PIP/PS lipid concentrations.
Membrane composition for live cell imaging is called PM (plasma membrane).
Cells were stimulated with 10 nM chemoattractant (fMLF).
* = assay buffer contains 75 mM NaCl instead of 150 mM NaCl.

#### PLASMID INVENTORY

| Recombinant DNA | | | |
|---|---|---|---|
| his6-TEV-SUMO3-GGGGG-PLCd PH domain (11-140aa) | bacterial | (ref) | pSH450 |
| his6-MBP-N10-TEV-GGGGG-Grp1 (261-387aa) | bacterial | This paper | pSH558 |
| his6-MBP-TEV-GGGG-TAPP1(182-303aa)-GGG-SNAP | bacterial | This paper | pSH1258 |
| his6-TEV-SUMO3-GGGGG-LactC2 (271-427aa) | bacterial | This paper | pSH1271 |
| his10-**mNG**-GGGGG-SHIP1 (PH-PP-C2, 292-878aa) | bacterial | This paper | pSH1082 |
| his6-TEV-SHIP1 FL (1-1188aa) | baculovirus | This paper | pSH798 |
| his6-TEV-**mNG**-SHIP1 FL (1-1188aa) | baculovirus | This paper | pSH973 |
| his6-TEV-**mNG**-SHIP1 ΔCT (1-878aa) | baculovirus | This paper | pSH1042 |
| his6-TEV-**mNG**-SHIP1 ΔSH2 (102-1188aa) | baculovirus | This paper | pSH1053 |
| psPAX2 (2nd generation lentiviral packaging vector) | lentivirus | Addgene, 12260 | pSH1224 |
| pVSV-G (VSV-G envelop protein) | lentivirus | Addgene, 138479 | pSH1226 |
| Ubc-**mEos3.2**-(GGGGS)x2-Grp1 (261-387aa) | lentivirus | This paper | pSH1252 |
| Ubc-**mEos3.2**-(GGGGS)x2-LactC2 (271-427aa) | lentivirus | This paper | pSH1267 |
| Ubc-**mEos3.2**-(GGGGS)x2-SHIP1 (PH-PP-C2, 292-878aa) | lentivirus | This paper | pSH1254 |
